# Bisantrene potentiates tyrosine kinase inhibitor activity in clear cell renal cell carcinoma

**DOI:** 10.1101/2025.07.21.665824

**Authors:** Joshua S. Brzozowski, Sumit Sahni, Heather C. Murray, Lauren Watt, Dylan Kiltschewskij, Murray J. Cairns, Marinella Messina, Daniel Tillett, Michael J. Kelso, Nicole M. Verrills

**Affiliations:** School of Biomedical Sciences and Pharmacy, College of Health, Medicine and Wellbeing, University of Newcastle, Newcastle, New South Wales, Australia and Precision Medicine Program, Hunter Medical Research Institute, New Lambton, New South Wales, Australia; Race Oncology Limited, Sydney, New South Wales, Australia

**Keywords:** Bisantrene, renal cell carcinoma, lenvatinib, pazopanib, cabozantinib, tyrosine kinase inhibitors

## Abstract

Clear Cell Renal Cell Carcinoma (ccRCC) is the most prevalent kidney cancer and often develops resistance to standard therapies. This study aimed to assess if bisantrene, a multi-mechanistic agent with broad anticancer activity, can enhance the activity of standard of care ccRCC treatments. A panel of ccRCC cell lines were treated with bisantrene alone and in combination with common ccRCC drugs. Bisantrene showed moderate activity as a single agent, but strongly synergized with several ccRCC treatments, especially the tyrosine kinase inhibitors (TKIs) lenvatinib, pazopanib and cabozantinib. Cellular signaling pathways assessed by immunoblotting revealed the TKIs inhibit MET as well as downstream AKT and ERK signaling pathways as single agents. Combination of these TKIs with bisantrene was able to induce sustained downstream AKT inhibition and negate the rebound effect seen with TKI resistance. Overall, bisantrene shows promise as a new therapeutic agent for ccRCC in combination with TKIs.

## 1. Introduction

Kidney cancer is the 12^th^ most common cancer worldwide ^1^, with renal cell carcinoma (RCC) accounting for 90% of all kidney cancers ^2, 3^. Risk factors for RCC include hypertension, obesity, smoking, age (> 60 years) and male sex ^1, 4^. RCCs are classified histologically into clear cell (ccRCC) and non-clear cell ^5^. ccRCC is the most common form (70-80%) and is often associated with mutations in the tumor suppressor gene, *Von Hippel-Lindau* (*VHL*) ^5-8^. Disruption of *VHL* increases the expression of Hypoxia Inducible Factors (HIFs), Vascular Endothelial Growth Factor (VEGF) and Platelet-Derived Growth Factor (PDGF), culminating in increased angiogenesis ^9, 10^. Inhibitors of the VEGF and PDGF tyrosine kinases and their downstream signaling pathways (e.g., mammalian target of rapamycin (mTOR)) are approved treatments for ccRCC ^11^. The tyrosine kinase inhibitors (TKIs) axitinib, cabozantinib, lenvatinib, pazopanib, sorafenib and sunitinib are routinely used as first or second line treatments in ccRCC ^12 2, 13-16^.

A key problem in ccRCC treatment is therapeutic resistance to TKI therapy. Temsirolimus and everolimus, both mTOR inhibitors, are indicated when treatment with TKIs fails ^17^. Immune checkpoint inhibitors are also used, either alone or together with TKIs ^18^. While these targeted treatments have improved the survival, new approaches that can overcome or prevent TKI resistance in ccRCC are required.

Bisantrene is an anthracene-based, anticancer drug that acts via several molecular mechanisms, including: binding and intercalation within the helices of deoxyribonucleic acid (DNA) ^19^, disruption of DNA replication ^20^, inhibiting angiogenesis ^21^, inhibiting telomerase ^22^, G-quadraplex binding ^23^, and inhibition of Topoisomerase II leading to double stranded DNA breaks ^24^. Bisantrene has been shown to inhibit both DNA and RNA synthesis ^20^ and induce c-Myc downregulation ^25^.

Bisantrene also displays immune-stimulatory effects that may contribute to its antitumour activity ^26,27^. Recently, bisantrene was reported to modulate the m^6^A RNA epigenetic pathway, which plays an important oncogenic role in various cancers by altering the activity of the m^6^A RNA demethylase Fatso/Fat Mass and Obesity-associated (FTO). ^25^ Inhibition of FTO has been shown to increase the global m^6^A levels in RNA in multiple cancer types ^25, 28-30^.

Early clinical investigations assessed bisantrene as a monotherapy in a range of cancers, including RCC, and some activity was noted ^31-36^. These historical studies support exploration of bisantrene in the modern RCC treatment context, especially in combination with more recent drugs. In this study, we investigated the use of bisantrene in combination with standard of care ccRCC TKIs and mTOR inhibitors using a panel of *VHL* mutant and wildtype ccRCC cell lines. Bisantrene exhibited moderate potency on its own, but showed unexpected and strong synergy with TKIs, most notably cabozantinib, lenvatinib, pazopanib, as well as the mTOR inhibitor, everolimus. These results support the potential of bisantrene use in combination with TKIs to improve treatment outcomes in ccRCC patients.

## 2. Methods

### 2.1 Drugs and cell lines

Bisantrene dihydrochloride (herein referred to as bisantrene; Race Oncology Ltd, Sydney, Australia) was reconstituted in dimethylsulphoxide (DMSO) at 20 mM. All other drugs were purchased from Selleck Chemicals (Houston, TX). Everolimus (RAD001), sunitinib, sorafenib, pazopanib, axitinib and cabozantinib (BMS-907351) were reconstituted in DMSO at 100 mM, lenvatinib at 17.2 mM, tivozanib at 50 mM and temsirolimus (CCI-779) at 80 mM. All drugs were stored at -20 °C as aliquots to reduce the number of freeze-thaw cycles.

Human ccRCC cell lines 769-P, 786-O, Caki-1, Caki-2, A-498 and A-704 were purchased from ATCC (Manassas, VA); RCC4-EV and RCC4-VHL from Millipore; and KMRC-1 from the Australian Cell Bank (New South Wales, Australia). The papillary RCC cell line, ACHN (ATCC), was included as an aggressive RCC model with wildtype *VHL* gene ^37^. All cells were cultured in a humidified chamber at 37 °C with 5% CO_2_ in the media listed in **Supplementary Table 1**. All human cell lines were authenticated using STR profiling within the last three years and experiments were performed with mycoplasma-free cells.

### 2.2 Cytotoxicity Assay

Cell viability was determined using a resazurin metabolic activity assay, as previously described ^38^. Briefly, cells were seeded in duplicate wells of 96-well microtiter plates at 1 x 10^3^ cells/well (786-O, RCC4 EV, RCC4 VHL, KMRC-1), 3 x 10^3^ cells/well (Caki-1, Caki-2, 769-P, A-498) or 5 x 10^3^ cells/well (A-704, ACHN) and cultured for 24 hours before addition of drugs and incubated for a further 72 hours. Viability was determined after addition of 1/10^th^ volume resazurin (0.6 mM resazurin, 78 μM methylene blue, 1 mM potassium hexacyanoferrate (III), 1 mM potassium hexacyanoferrate (II) trihydrate (Merck; Victoria, Australia) in phosphate buffered saline (PBS)) for 5 hours and fluorescence measured at 544 nm excitation/590 nm emission on a FLUOstar OPTIMA plate reader (BMG Lab Technologies; Ortenberg, Germany). Graphpad Prism 9 software (Boston, MA) was used to generate graphs. Drug IC_50_ values were determined by cubic spline-lowess regression analysis using Graphpad Prism 9. At least three independent replicates were performed for each cell line and each drug combination, and data are represented as mean ± standard error of the mean (SEM).

### 2.3 Clonogenicity Assay

Clonogenicity assays were used to determine the colony-forming ability of cells treated with bisantrene. Cells were seeded into 6-well plates at 150 cells/well (786-O), 1000 cells/well (RCC4 EV, RCC4 VHL, A-498) or 2000 cells/well (A-704, 769-P, Caki-1, Caki-2, KMRC-1, ACHN) and allowed to adhere for 24 hours before addition of bisantrene and incubation for 4 days. Drug-containing media was removed and replaced with fresh media (without drug) and cultured for an additional 4 days prior to fixing with ice-cold methanol for 10 minutes at 4 °C and staining with crystal violet solution (0.5% crystal violet, 25% methanol in PBS). Colonies were captured on a ChemiDoc MP Imaging System (Bio-Rad). Images were analyzed using the ColonyArea plugin ^39^ for ImageJ ^40^ and the percent area of the well filled by colonies was determined and presented as percentage of untreated cells using Graphpad Prism 9 software. Four independent replicates were performed for each cell line and data are represented as mean ± SEM.

### 2.4 Synergy Analysis

Synergy analysis for the combination drug treatments was conducted using the Bliss synergy method within SynergyFinder 2.0 software ^41, 42^. When the Bliss synergy score is less than -10, the interaction between two drugs is likely to be antagonistic. When the synergy score is from -10 to 10, the interaction is likely to be additive, and when the score is greater than 10, the interaction is likely to be synergistic. The fractional product method of Webb was utilized as a second method for verifying synergy between the drugs ^43^. Webb analyses are interpreted as: values < -0.1 indicate synergy, between -0.1 and 0.1 indicate additivity and values > 0.1 indicate antagonism.

### 2.5 Immunoblotting

Whole-cell lysates were sonicated 2 x 10s in ice-cold radioimmunoprecipitation assay buffer freshly supplemented with protease inhibitors (0.05 M HEPES [pH 7.4], 1% Triton X-100, 0.1% SDS, 50 mM sodium fluoride, 0.05 M EDTA, 5% sodium deoxycholate, 1 mM sodium orthovanadate, and Protease Inhibitor Cocktail [Merck]). Total protein was quantified by bicinchoninic acid assay, as per the manufacturer’s instructions (Thermo Fisher Scientific; New South Wales, Australia). Lysates were resolved in reducing conditions on 4 to 12% gradient Bis–Tris NuPAGE precast gels (Thermo Fisher Scientific) before wet transfer to nitrocellulose. The LI-COR Odyssey M imaging system (LI-COR; Lincoln, NE) with IRDye conjugated antibodies was used to probe blots with both phosphorylated and total protein. All primary antibodies used were from Cell Signaling Technology: p-Akt Thr308 (Cat#9275), Akt (Cat#2920), pERK1/2 T202/Y204 (mouse T203/Y205, Cat#9101S), ERK1/2 (Cat#9107S), p-Met Tyr1234/1235 (Cat#3077) and Met (Cat#8198), except GAPDH, which was purchased from Biovision (Milpitas, CA; Cat#3777R) as a loading control. Secondary antibodies were IRDye 800CW anti-mouse or IRDye 680RD anti-rabbit secondary antibody (LI-COR). Densitometry was performed using Empiria Studio software (LI-COR).

### 2.6 Data Analysis

Data were statistically compared using either an unpaired Student’s *t*-test when comparing two groups, or one-way ANOVA followed by Tukey’s post-hoc test for multiple comparisons. Statistical analysis was performed using GraphPad Prism software. Statistically significant differences were observed at *p < 0.05*. At least three independent replicates were performed for each experiment. Data are presented as mean ± SEM.

## 3. Results

### 3.1 Activity of bisantrene in ccRCC cell lines

To assess the sensitivity of ccRCC cell lines to bisantrene as a single agent, cells were treated using a range of bisantrene concentrations for 72 h and cell viability determined using a resazurin metabolic assay. The drug concentration required to inhibit cell viability by 50% (IC_50_) was determined for each cell line (**Figure 1A**; **Table 1**). Apart from the A-704 cell line, which was highly resistant to bisantrene (IC_50_: 12.3 μM), all ccRCC cells showed IC_50_ values below 1.31 μM, with 5 of the 9 lines showing IC_50_ below 1 μM. The resistance of A-704 cells to bisantrene may be due to their low tumorigenic nature, as suggested by these cells not forming tumors in immunosuppressed mice, while they do form colonies in semisolid media ^44^. Bisantrene also showed potent activity against the papillary RCC cell line, ACHN (IC_50_: 242 nM).

**Figure 1.**
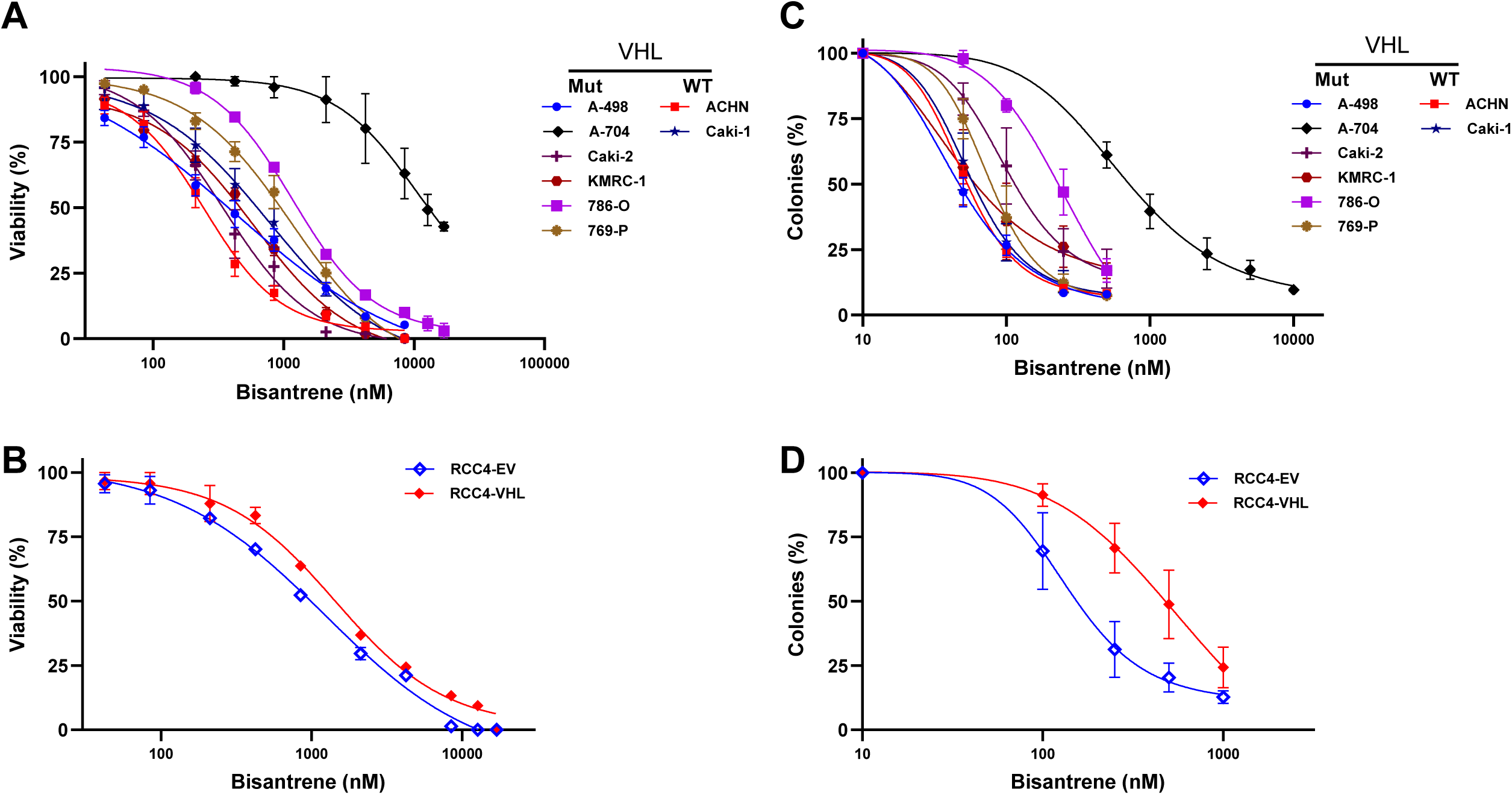
Effect of bisantrene on viability and clonogenicity of RCC cell lines (A,B) Cell Viability Assay: Cells were treated with bisantrene for 72 h using the indicated drug concentrations and cell viability determined using a resazurin metabolic assay. **(A)** Cell viability is expressed as a percentage of untreated control cells. **(B)** Cytotoxicity of bisantrene was compared in the parental RCC4-EV (mutant VHL) versus the RCC4-VHL (WT VHL rescue) cell lines. **(C,D) Clonogenicity Assay:** Cells were treated with bisantrene for 96 h using the indicated drug concentrations then fresh drug-free media was added for a further 96 h. Colonies were detected using crystal violet and quantified using Image J. Colonies are expressed as a percentage of untreated control cells. **(C)** Clonogenic survival dose response for cell lines tested. **(D)** Clonogenic survival dose response for the RCC4-EV and RCC4-VHL (WT VHL rescue) cell lines. n ≥ 3.

**Table 1:**
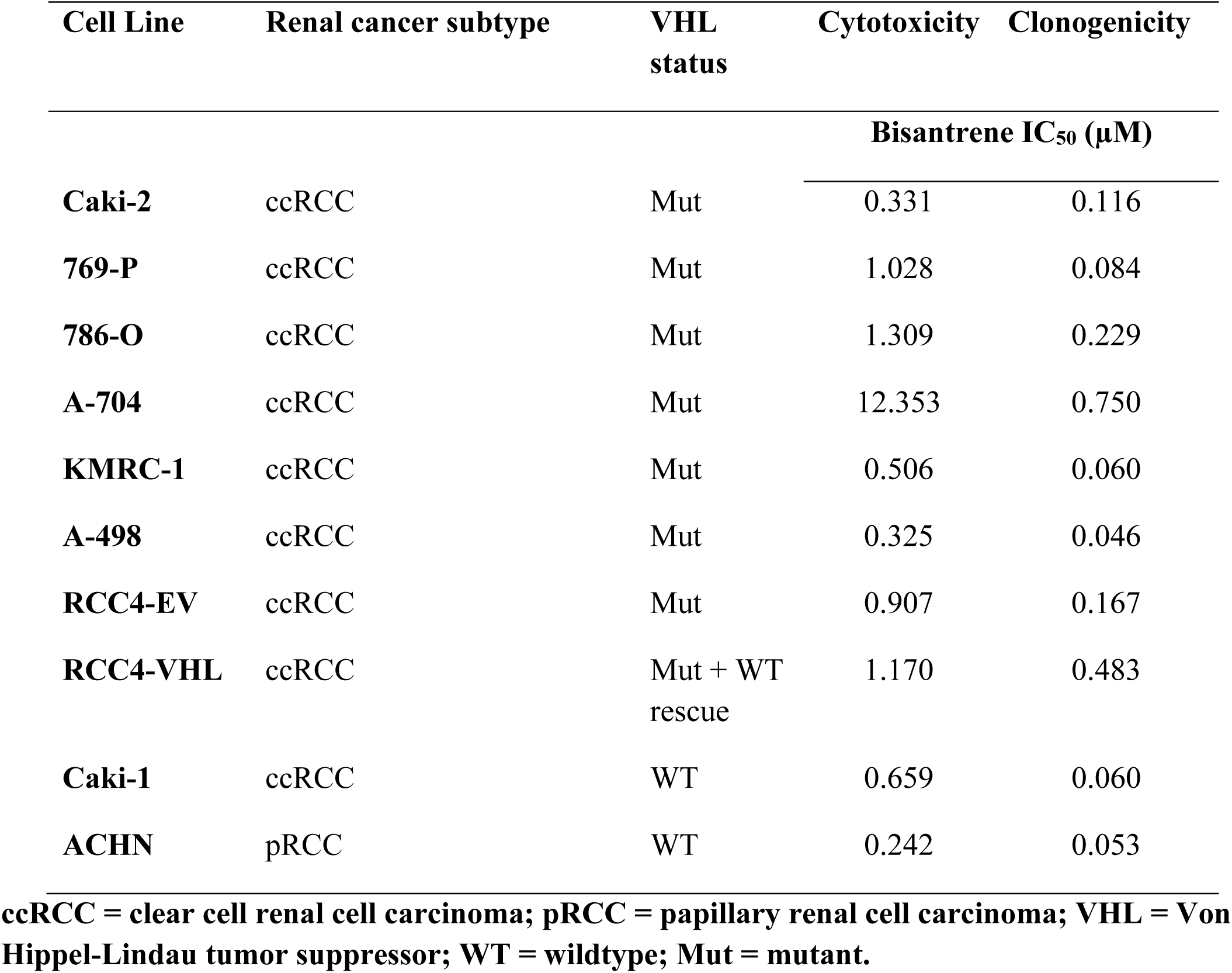
Growth inhibitory effects of bisantrene in RCC (ccRCC and pRCC) cells.

| Cell Line | Renal cancer subtype | VHL status | Cytotoxicity | Clonogenicity |
| --- | --- | --- | --- | --- |
| Bisantrene IC<sub>50</sub> (μM) |  |  |  |  |
| **Caki-2** | ccRCC | Mut | 0.331 | 0.116 |
| **769-P** | ccRCC | Mut | 1.028 | 0.084 |
| **786-O** | ccRCC | Mut | 1.309 | 0.229 |
| **A-704** | ccRCC | Mut | 12.353 | 0.750 |
| **KMRC-1** | ccRCC | Mut | 0.506 | 0.060 |
| **A-498** | ccRCC | Mut | 0.325 | 0.046 |
| **RCC4-EV** | ccRCC | Mut | 0.907 | 0.167 |
| **RCC4-VHL** | ccRCC | Mut + WT rescue | 1.170 | 0.483 |
| **Caki-1** | ccRCC | WT | 0.659 | 0.060 |
| **ACHN** | pRCC | WT | 0.242 | 0.053 |

ccRCC = clear cell renal cell carcinoma; pRCC = papillary renal cell carcinoma; VHL = Von Hippel-Lindau tumor suppressor; WT = wildtype; Mut = mutant.

Comparison of the RCC4-EV line carrying an endogenous mutant *VHL* and the isogenic ‘rescue’ cell line, RCC4-VHL, which has been transduced with the wildtype *VHL* gene to rescue *VHL* loss, revealed that the mutant was slightly more sensitive (1.3-fold, *p < 0.05*) to bisantrene than the WT-VHL rescue cell line (**Figure 1B**).

### 3.2 Effect of bisantrene on colony formation in ccRCC cell lines

The effects of bisantrene were also assessed using long term colony formation (clonogenicity) assays. All cell lines were more sensitive to bisantrene in this assay compared to the cytotoxicity assays (**Table 1**). Similar sensitivity trends were observed for the cell lines across both assays, with the A-704 cells being the most resistant to bisantrene, and the ACHN, KMRC-1 and A-498 cell lines being the most sensitive (**Figure 1C**). Notably, the Caki-1 cells exhibited markedly greater sensitivity to bisantrene in the clonogenic assays compared to cytotoxicity assays. Similar to the cytotoxicity assay, the RCC4-EV cells were significantly more sensitive (2.9-fold, *p* < 0.05) to bisantrene than the RCC4-VHL WT rescue cells in the clonogenic assay (**Figure 1D**; **Table 1**).

### 3.3 Synergistic cytotoxicity of bisantrene in combination with ccRCC drugs

To investigate the activity of bisantrene when co-administered with current ccRCC drugs, combination cytotoxicity assays were performed using the 786-O, Caki-1, Caki-2, RCC4-EV and RCC4-VHL cell lines. Single agent assays were first performed to determine the IC_50_ of lenvatinib, pazopanib, cabozantinib, sunitinib, sorafenib, axitinib, temsirolimus and everolimus (**Table 2, Supplementary Figure 1**). Each drug was then tested in combination with bisantrene at 5 different concentrations encompassing the IC_50_ for each drug. The effects of drug combination concentrations on cell viability are presented as red/blue heatmaps (**Figure 2; Supplementary Figure 2**). The fractional product method of Webb was used to determine whether the effects of the combined drug treatments were additive, synergistic or antagonistic ^43^. Synergy was observed with sunitinib, sorafenib, axitinib, everolimus and temsirolimus, most frequently at lower concentrations of bisantrene (**Supplementary Figure 2**). The greatest synergy was observed for bisantrene in combination with the kinase inhibitors lenvatinib (**Figure 2A**), pazopanib (**Figure 2B**) and cabozantinib (**Figure 2C**).

**Figure 2.**
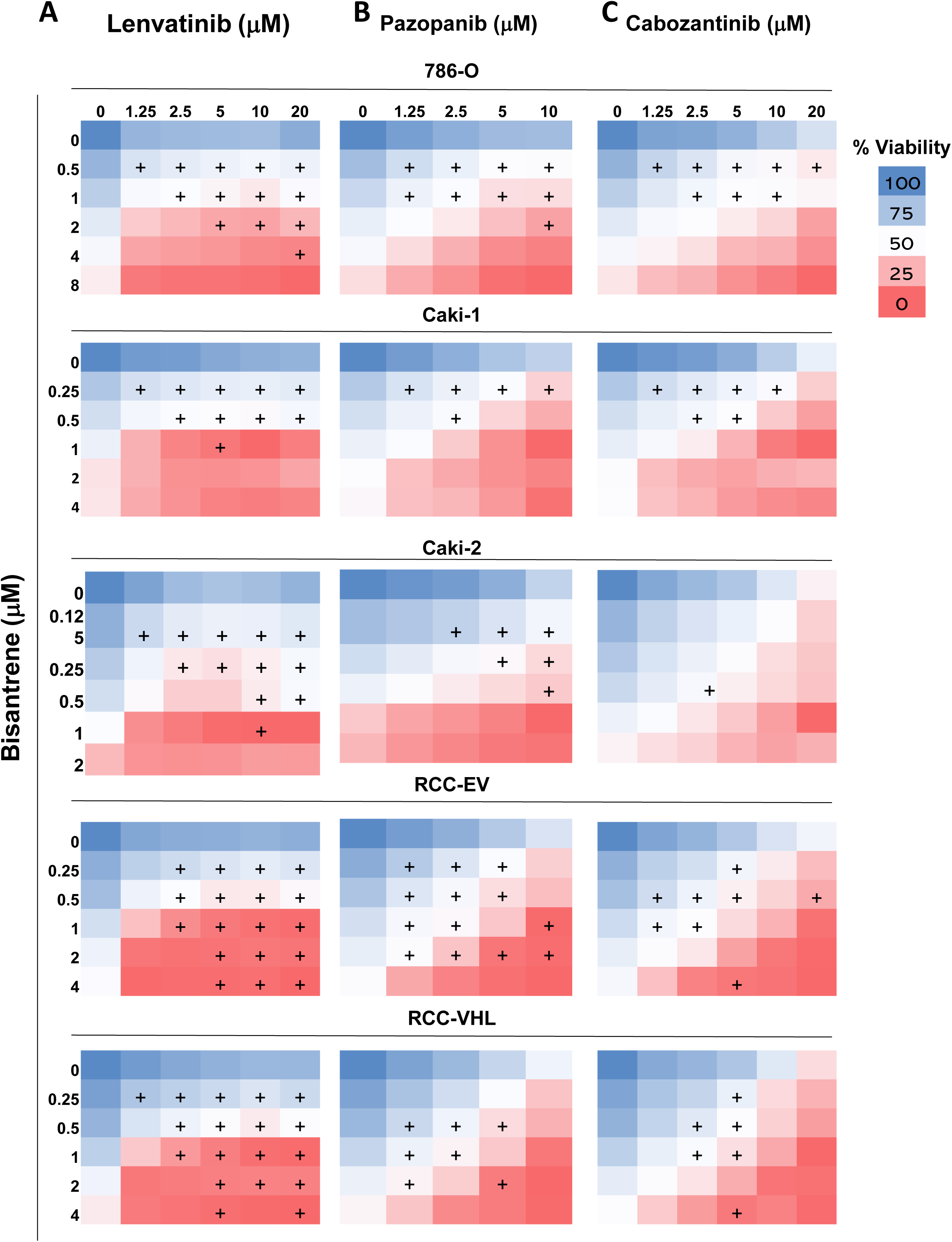
Cell viability in response to combined drug treatment with bisantrene and current ccRCC drugs in ccRCC cell lines. Different ccRCC cell lines (786-O, Caki-1, Caki-2, RCC-EV and RCC-VHL) were treated with the indicated concentrations of bisantrene and ccRCC drugs for 72 h and cell viability determined using a resazurin assay. ccRCC drugs used: **(A)** lenvatinib**; (B)** pazopanib; and **(C)** cabozantinib. Average percent viability relative to untreated cells is presented as a heat map, where blue indicates high viability and red indicates low viability. + signs indicate synergy, as assessed by the fractional product method of Webb ^43^.

**Table 2:** IC_50_ of single agent ccRCC drugs used in combination cytotoxicity assays.

| Cell line | IC<sub>50</sub> (μM) |  |  |  |  |  |  |  |
| --- | --- | --- | --- | --- | --- | --- | --- | --- |
|  | Len | Paz | Cabo | Suni | Soraf | Axit | Tem | Ever |
| **786-O** | >20 | >10 | 11.728 | 3.702 | 7.916 | 2.479 | 0.129 | 1.467 |
| **Caki-1** | >20 | 7.867 | 10.442 | >8 | 9.658 | >8 | >2 | >20 |
| **Caki-2** | >20 | >10 | 7.345 | 5.451 | 15.997 | >8 | >2 | >20 |
| **RCC4-EV** | >20 | 7.021 | 6.586 | 5.02 | 10.233 | >8 | 0.109 | 1.119 |
| **RCC4-VHL** | >20 | 5.202 | 7.326 | 2.216 | 8.65 | 4.0575 | 0.124 | 1.151 |

Len: Lenvatinib; Paz: Pazopanib; Cabo: Cabozantinib; Suni: Sunitinib; Soraf: Sorafenib; Axit: Axitinib; Tem: Temsirolimus; Ever: Everolimus.

To verify the synergy results, a second analysis was performed using the Bliss method (**Table 3; Supplementary Figures 3**) ^42^. This analysis revealed synergistic effects for bisantrene with lenvatinib and pazopanib in all five cell lines, while bisantrene and cabozantinib were synergistic in all cells except Caki-2, where an additive effect was seen (**Table 3**). Temsirolimus showed synergy with bisantrene in the Caki-1, Caki-2 and RCC4-EV cell lines, with some synergy observed with sunitinib or sorafenib in the 786-O cells, and axitinib in the Caki-1 cells (**Table 3**). In general, RCC4-EV cells with mutant *VHL* showed higher overall Bliss synergy scores for bisantrene combinations with TKIs compared to wildtype rescue RCC4-VHL cells (**Table 3**). The most synergistic area across drug doses determined by Bliss analysis further suggested that areas of synergy were present for almost all drugs assessed across all cell lines, except for axitinib (**Table 3**). Lenvatinib, pazopanib and cabozantinib had the highest scores for the most synergistic area. Individual Bliss scores for each concentration combination further revealed multiple synergistic combinations across most cell lines (**Supplementary Table 2**). After identifying that several TKIs demonstrated high synergy with bisantrene, we chose to evaluate tivozanib, a recently approved TKI for ccRCC ^45^. As observed with the other TKIs, tivozanib showed potent synergy with bisantrene in 786-O ccRCC cells (**Supplementary Figure 4**).

**Table 3.** Synergy scores for the combination of bisantrene and current ccRCC drugs, as determined by Bliss analysis in different ccRCC cell lines ^42^.

| Drug combination | Cell Line |  |  |  |  |  |  |  |  |  |
| --- | --- | --- | --- | --- | --- | --- | --- | --- | --- | --- |
|  | 786-0 |  | Caki-1 |  | Caki-2 |  | RCC4-EV |  | RCC4-VHL |  |
| Bisantrene + | OSS<sup>1</sup> | MSAS<sup>2</sup> | OSS | MSAS | OSS | MSAS | OSS | MSAS | OSS | MSAS |
| Lenvatinib | 24.49 | 35.02 | 20.69 | 30.28 | 17.93 | 21.24 | 36.42 | 44.15 | 29.32 | 36.05 |
| Pazopanib | 21.86 | 30.44 | 16.24 | 22.50 | 13.95 | 16.90 | 22.34 | 26.02 | 17.84 | 22.96 |
| Cabozantinib | 18.07 | 30.13 | 15.08 | 26.66 | 8.64 | 18.54 | 19.15 | 26.86 | 15.21 | 24.50 |
| Sunitinib | 12.26 | 24.37 | 6.42 | 14.24 | 5.26 | 15.73 | 8.01 | 14.79 | 5.21 | 12.97 |
| Sorafenib | 10.40 | 23.69 | 7.86 | 18.66 | 5.07 | 11.54 | 5.09 | 10.41 | 4.41 | 12.14 |
| Axitinib | 4.25 | 9.19 | 10.51 | 15.29 | 1.07 | 3.59 | 5.15 | 11.68 | 0.09 | 6.90 |
| Temsirolimus | 8.84 | 14.76 | 15.69 | 23.26 | 10.34 | 17.49 | 10.27 | 15.85 | 8.43 | 18.58 |
| Everolimus | 18.15 | 27.12 | 19.88 | 28.81 | 15.45 | 23.00 | 17.58 | 22.31 | 17.74 | 27.52 |

<sup>1</sup> OSS, Overall synergy score. <sup>2</sup> MSAS, Most synergistic area score. Values >10 indicate synergy (bold); values between -10 to 10 are additive; values below -10 indicate antagonism.

### 3.4 Combining bisantrene with TKIs induces sustained inhibition of the AKT pathway

We explored the potential mechanisms behind bisantrene TKI synergy with lenvatinib and cabozantinib. Both TKIs are known to inhibit multiple kinases, including VEGFR, with cabozantinib also inhibiting MET, the receptor for hepatocyte growth factor ^46, 47^. As a consequence, we focused on total and activated (phosphorylated) MET, as well as the key convergence points for tyrosine kinase receptor signaling AKT and ERK after immediate (1 h) and prolonged (24 h) drug treatments in 786-O, Caki-1 and Caki-2 cells.

Bisantrene as a single agent inhibited activation (phosphorylation) of MET only after prolonged (24 h) treatment. In contrast, both TKIs were able to significantly reduce pMET after short treatments and the effect remained after 24 h **(Figure 3**). Combined treatment with bisantrene and TKIs did not further reduce the amount of pMET at either timepoint relative to TKI alone. (**Figure 3**).

**Figure 3.**
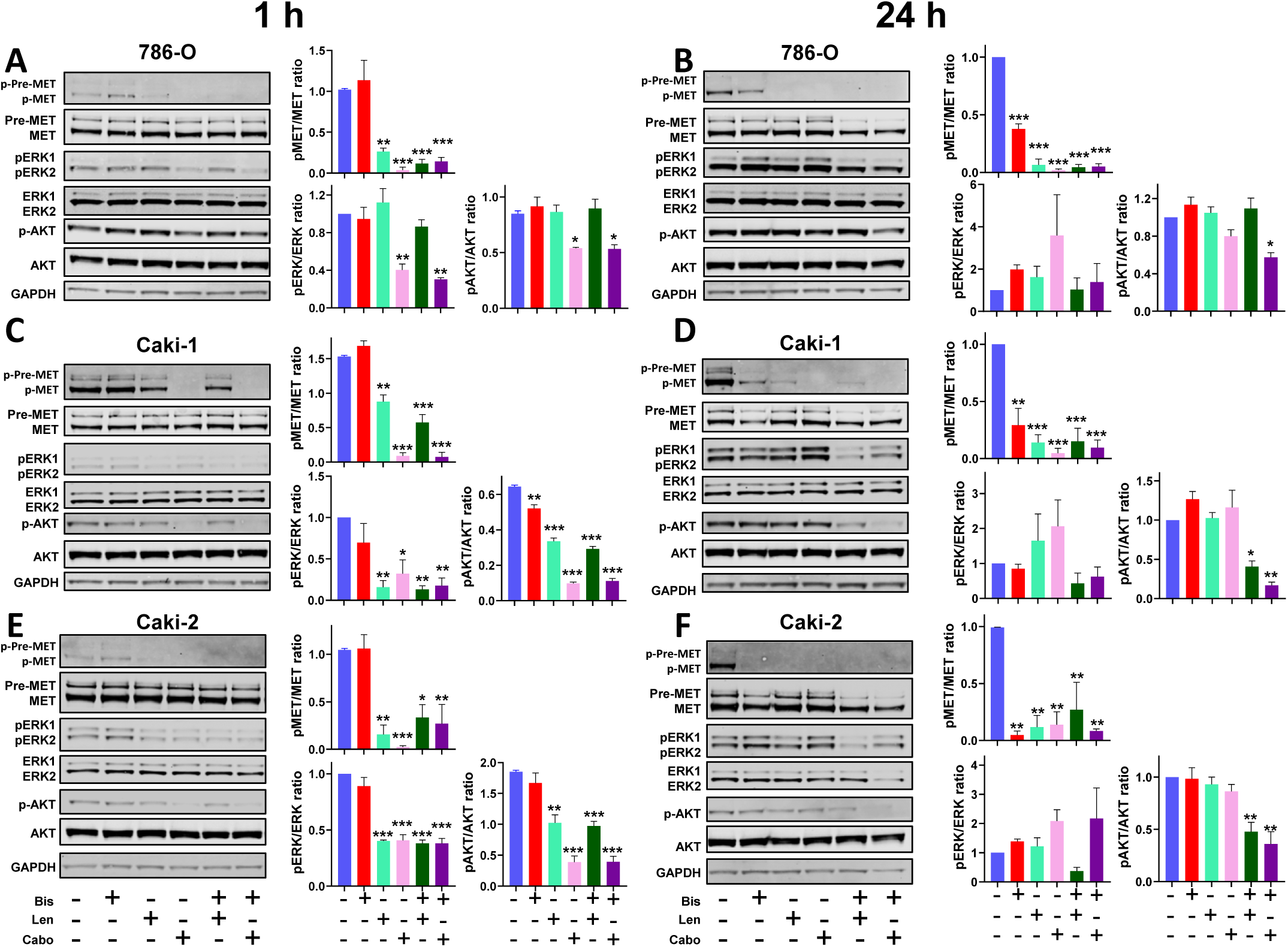
Cellular signaling in response to treatment with bisantrene and TKIs in 786-O, Caki-1 and Caki-2 ccRCC cells. Cells were treated with 2 μM bisantrene (Bis), 10 μM lenvatinib (Len), 10 μM cabozantinib (Cabo) or drug combinations as shown for 1 h (**A, C, E**) or 24 h (**B, D, F**) prior to western blotting of total cell lysates probed with antibodies to indicated proteins. Phosphorylated and total proteins were imaged on the same blot using two different IRDyes. Images are representative of n = 3. Densitometric quantitation of each protein was performed and ratio of phosphorylated to total protein was determined and compared among different treatment groups. *\*,p<0.05, **p<0.01, ***p<0.001* compared to control.

Treatment with bisantrene alone had no effect on ERK activation (phosphorylation) in any cell line at either time point (**Figure 3; Supplementary Figure 5**). Cabozantinib alone suppressed the activation of ERK in 786-O, Caki-1 and Caki-2 cells after short-term (1 h) incubation, while lenvatinib reduced pERK in the Caki-1 and Caki-2 line, but not 786-O. There was no additional suppression of pERK observed at 1 h for either agent in these three cell lines when combined with bisantrene (**Figure 3**). After 24 h of drug exposure, a trend towards increased ERK activation was observed after TKI treatment compared to untreated cells, while treatment in combination with bisantrene appeared to suppress this activation, although these effects did not achieve statistical significance (**Figure 3**).

There was no significant effect of bisantrene alone on AKT activation (phosphorylation), with only a slight reduction observed following short-term treatment in Caki-1 cells (**Figure 3**). In contrast, a significant inhibition of AKT activation was observed after 1 h with both TKIs across all cells, except for lenvatinib in the 786-O line (**Figure 3**). No further reductions in pAKT were observed after 1 hour when bisantrene was added to the TKIs. Notably, after 24 h treatment, while none of the single agents were unable to suppress AKT activation, the combination of bisantrene with TKIs provided sustained inhibition of AKT in 786-O (cabozantinib + bisantrene, only) and Caki-1/Caki-2 cells (both TKIs + bisantrene) (**Figure 3**).

## 4. Discussion

Bisantrene is a multi-mechanistic anticancer agent with low cardiotoxicity ^48^. Early clinical studies suggested bisantrene may have single agent efficacy in treating RCC ^33, 35, 36^, but whether bisantrene could enhance the efficacy of modern targeted therapies for ccRCC was unknown. In this study, we demonstrated that bisantrene provides synergistic cytotoxicity when combined with approved TKIs, and to a lesser extent mTOR inhibitors, in ccRCC cell lines, supporting incorporation of bisantrene into TKI-based therapeutic strategies for ccRCC (**Figure 4**). An investigation into the possible mechanisms contributing to the synergy of bisantrene and TKIs suggested that sustained inhibition of the AKT signaling pathway may be involved (**Figure 4**).

**Figure 4.**
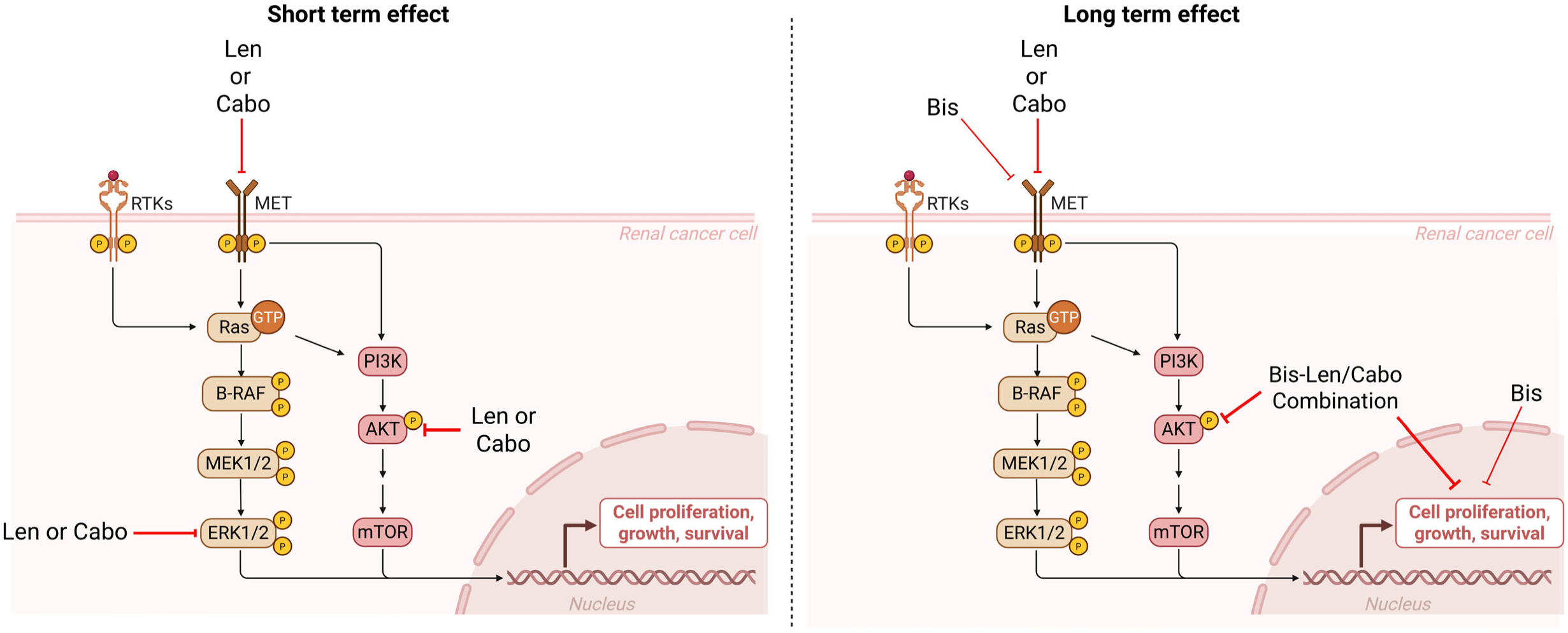
Schematic overview of observed synergistic activity between bisantrene and TKIs in ccRCC models after short-term and long-term treatments. Bis = Bisantrene; Len = Lenvatinib; Cabo = Cabozantinib.

TKIs are currently approved as first line treatments for advanced ccRCC patients ^49^, but development of resistance to these targeted therapies is inevitable ^9^. Use of a broad-spectrum, multi-mechanistic agent like bisantrene in combination with TKI-targeted therapy may offer a solution for overcoming or delaying resistance and addressing this significant clinical challenge. Bisantrene has multiple overlapping mechanisms of action with anthracyclines, including the ability to inhibit topoisomerase II ^48^. Unlike anthracyclines, bisantrene is not associated with significant cardiotoxicity risks ^48^, and therefore represents a promising broad-spectrum alternative for use in combination with TKIs for the treatment of ccRCC. Similar to previously reported effects of anthracyclines ^50, 51^, we found that bisantrene is significantly more active against *VHL* mutant cells (RCC4-EV) compared to WT *VHL* rescue cells (RCC4-VHL). Moreover, we observed a trend towards higher synergy between bisantrene and TKIs in RCC4-EV cells compared to RCC4-VHL cells. Given that *VHL* mutations are present in up to 91% of ccRCC cases ^52^, these results support the potential of bisantrene in combination with TKIs as a possible strategy in a significant subset of ccRCC patients. VHL acts as an E3 ubiquitin ligase to promote proteasomal degradation of HIF proteins, but is also known to play a role in other cellular processes ^52^. VHL has been reported to suppress transcription of the *MYC* oncogene by binding to its promoter region ^53^. Interestingly, bisantrene is also able to downregulate the expression of *MYC* ^25^, which may explain the increased sensitivity of *VHL* mutant ccRCC cells to bisantrene.

Upregulation of the AKT signaling pathway is a driver of TKI resistance in ccRCC ^9^. We observed that while TKIs initially suppress AKT activation after short-term exposure, this effect was lost after prolonged treatment. In general, the combination of bisantrene with TKIs was able to suppress the AKT pathway for prolonged periods, suggesting this as a potential mechanism for the synergy seen with bisantrene and TKIs in ccRCC.

## 5. Conclusions

Bisantrene demonstrates single agent cytotoxicity against ccRCC cells and shows strong synergy with several currently approved ccRCC drugs, particularly the TKIs lenvatinib, cabozantinib and pazopanib. While bisantrene, lenvatinib and cabozantinib are all able to inhibit activation of the tyrosine kinase MET, only combinations of bisantrene with lenvatinib or cabozantinib can provide sustained suppression of AKT signaling, a common resistance pathway observed in receptor tyrosine kinases involved in ccRCC. Overall, these results support further investigation of combinations of bisantrene with TKIs as a strategy for overcoming TKI resistance in ccRCC.

## List of abbreviations

ccRCC: Clear Cell Renal Cell Carcinoma
PDGF: Platelet-Derived Growth Factor
mTOR: Mammalian Target of Rapamycin
Mut: Mutant
RCC: Renal Cell Carcinoma
TKI: Tyrosine Kinase Inhibitor
VEGF: Vascular Endothelial Growth Factor
VEGFR: Vascular Endothelial Growth Factor Receptor
VHL: Von Hippel-Lindau
WT: Wildtype

## Availability of data and materials

The datasets used and/or analyzed during the current study are available from the corresponding author on reasonable request.

## Author information

JB conducted all cell experiments; JB, SS, HCM, LW, DK and NMV performed data analysis and figure generation; SS, LW and NMV prepared the manuscript. MM, DT, MJK, MC and NMV contributed to project design and interpretation. NMV provided project management.

## References

1 Cirillo L, Innocenti S, Becherucci F. Global epidemiology of kidney cancer. Nephrology Dialysis Transplantation 2024; 39 (6): 920–928.

2 Hsieh JJ, Purdue MP, Signoretti S et al. Renal cell carcinoma. Nature Reviews Disease Primers 2017; 3 (1): 17009.

3 Motzer RJ, Jonasch E, Agarwal N, et al. Kidney Cancer, Version 2.2017, NCCN Clinical Practice Guidelines in Oncology. J Natl Compr Canc Netw 2017; 15 (6): 804–834.

4 Capitanio U, Bensalah K, Bex A et al. Epidemiology of Renal Cell Carcinoma. Eur Urol 2019; 75 (1): 74–84.

5 Muglia VF, Prando A. Renal cell carcinoma: histological classification and correlation with imaging findings. Radiol Bras 2015; 48 (3): 166–174.

6 Haas NB, Nathanson KL. Hereditary kidney cancer syndromes. Adv Chronic Kidney Dis 2014; 21 (1): 81–90.

7 Maher ER. Hereditary renal cell carcinoma syndromes: diagnosis, surveillance and management. World Journal of Urology 2018; 36 (12): 1891–1898.

8 Ricketts CJ, De Cubas AA, Fan H et al. The Cancer Genome Atlas Comprehensive Molecular Characterization of Renal Cell Carcinoma. Cell Rep 2018; 23 (1): 313–326.e315.

9 Duran I, Lambea J, Maroto P et al. Resistance to Targeted Therapies in Renal Cancer: The Importance of Changing the Mechanism of Action. Target Oncol 2017; 12 (1): 19–35.

10 Lai Y, Zhao Z, Zeng T et al. Crosstalk between VEGFR and other receptor tyrosine kinases for TKI therapy of metastatic renal cell carcinoma. Cancer Cell Int 2018; 18: 31.

11 Sheng X, Ye D, Zhou A et al. Efficacy and safety of vorolanib plus everolimus in metastatic renal cell carcinoma: A three-arm, randomised, double-blind, multicentre phase III study (CONCEPT). Eur J Cancer 2023; 178: 205–215.

12 Bahadoram S, Davoodi M, Hassanzadeh S et al. Renal cell carcinoma: an overview of the epidemiology, diagnosis, and treatment. G Ital Nefrol 2022; 39 (3): 2022-vol2023.

13 Escudier B, Eisen T, Stadler WM et al. Sorafenib in advanced clear-cell renal-cell carcinoma. N Engl J Med 2007; 356 (2): 125–134.

14 Motzer RJ, McCann L, Deen K. Pazopanib versus sunitinib in renal cancer. N Engl J Med 2013; 369 (20): 1970.

15 Rini BI, Escudier B, Tomczak P et al. Comparative effectiveness of axitinib versus sorafenib in advanced renal cell carcinoma (AXIS): a randomised phase 3 trial. Lancet 2011; 378 (9807): 1931–1939.

16 Choueiri TK, Escudier B, Powles T et al. Cabozantinib versus Everolimus in Advanced Renal-Cell Carcinoma. New England Journal of Medicine 2015; 373 (19): 1814–1823.

17 Patel SB, Stenehjem DD, Gill DM et al. Everolimus Versus Temsirolimus in Metastatic Renal Cell Carcinoma After Progression With Previous Systemic Therapies. Clin Genitourin Cancer 2016; 14 (2): 153–159.

18 Makhov P, Joshi S, Ghatalia P et al. Resistance to Systemic Therapies in Clear Cell Renal Cell Carcinoma: Mechanisms and Management Strategies. Mol Cancer Ther 2018; 17 (7): 1355–1364.

19 Wunz TP, Craven MT, Karol MD et al. DNA binding by antitumor anthracene derivatives. Journal of Medicinal Chemistry 1990; 33 (6): 1549–1553.

20 Citarella RV, Wallace RE, Murdock KC et al. Activity of a novel anthracenyl bishydrazone, 9,10-anthracenedicarboxyaldehyde Bis[(4,5-dihydro-1H-imidazol-2-yl)hydrazone] dihydrochloride, against experimental tumors in mice. Cancer Res 1982; 42 (2): 440-444.

21 Polverini PJ, Novak RF. Inhibition of angiogenesis by the antineoplastic agents mitoxantrone and bisantrene. Biochemical and Biophysical Research Communications 1986; 140 (3): 901–907.

22 Perry PJ, Reszka AP, Wood AA et al. Human Telomerase Inhibition by Regioisomeric Disubstituted Amidoanthracene-9,10-diones. Journal of Medicinal Chemistry 1998; 41 (24): 4873–4884.

23 Folini M, Pivetta C, Zagotto G et al. Remarkable interference with telomeric function by a G-quadruplex selective bisantrene regioisomer. Biochemical Pharmacology 2010; 79 (12): 1781–1790.

24 Zagotto G, Oliva A, Guano F, et al. Synthesis, DNA-damaging and cytotoxic properties of novel topoisomerase II-directed bisantrene analogues. Bioorganic & Medicinal Chemistry Letters 1998; 8 (2): 121–126.

25 Su R, Dong L, Li Y et al. Targeting FTO Suppresses Cancer Stem Cell Maintenance and Immune Evasion. Cancer Cell 2020; 38 (1): 79–96.e11.

26 Wang BS, Lumanglas AL, Ruszala-Mallon VM, Durr FE. Activation of Tumor-cytostatic Macrophages with the Antitumor Agent 9,10-Anthracenedicarboxaldehyde Bis[(4,5-dihydro-1H-imidazole-2-yl)hydrazone] Dihydrochloride (Bisantrene). Cancer Research 1984; 44 (6): 2363–2367.

27 Wang BS, Lumanglas AL, Durr FE. Immunotherapy of a Murine Lymphoma by Adoptive Transfer of Syngeneic Macrophages Activated with Bisantrene. Cancer Research 1986; 46 (2): 503–506.

28 Cui Y-H, Yang S, Wei J et al. Autophagy of the m6A mRNA demethylase FTO is impaired by low-level arsenic exposure to promote tumorigenesis. Nature Communications 2021; 12 (1): 2183.

29 Garg R, Melstrom L, Chen J et al. Targeting FTO Suppresses Pancreatic Carcinogenesis via Regulating Stem Cell Maintenance and EMT Pathway. Cancers 2022; 14 (23): 5919.

30 Phan T, Nguyen VH, Su R et al. Targeting fat mass and obesity-associated protein mitigates human colorectal cancer growth in vitro and in a murine model. Front Oncol 2023; 13: 1087644.

31 Spadea A, Petti MC, Aloespiriti MA et al. Bisantrene in relapsed and refractory acute myelogenous leukemia. Leuk Lymphoma 1993; 9 (3): 217–220.

32 Cowan JD, Neidhart J, McClure S et al. Randomized trial of doxorubicin, bisantrene, and mitoxantrone in advanced breast cancer: a Southwest Oncology Group study. J Natl Cancer Inst 1991; 83 (15): 1077–1084.

33 Evans WK, Shepard FA, Blackstein ME et al. Phase II evaluation of bisantrene in patients with advance renal cell carcinoma. Cancer Treatment Reports 1984; 69 (5): 727–728.

34 Holmes FA, Esparza L, Yap HY et al. A comparative study of bisantrene given by two dose schedules in patients with metastatic breast cancer. Cancer Chemother Pharmacol 1986; 18 (2): 157–161.

35 Spiegel RJ, Levin M, Blum R. Phase II trial of daily x5 bisantrene in renal cell carcinoma. Proceedings of the American Association for Cancer Research 1982; 23 (435): 111.

36 Myers JW, von Hoff DD, Coltman CA, Jr. et al. Phase II evaluation of bisantrene in patients with renal cell carcinoma. Cancer Treatment Reports 1982; 66 (10): 1869–1871.

37 Wolf MM, Kimryn Rathmell W, Beckermann KE. Modeling clear cell renal cell carcinoma and therapeutic implications. Oncogene 2020; 39 (17): 3413–3426.

38 Roberts KG, Smith AM, McDougall F et al. Essential requirement for PP2A inhibition by the oncogenic receptor c-KIT suggests PP2A reactivation as a strategy to treat c-KIT+ cancers. Cancer Res 2010; 70 (13): 5438–5447.

39 Guzmán C, Bagga M, Kaur A et al. ColonyArea: an ImageJ plugin to automatically quantify colony formation in clonogenic assays. PLoS One 2014; 9 (3): e92444.

40 Schneider CA, Rasband WS, Eliceiri KW. NIH Image to ImageJ: 25 years of image analysis. Nat Methods 2012; 9 (7): 671–675.

41 Ianevski A, Giri AK, Aittokallio T. SynergyFinder 2.0: visual analytics of multi-drug combination synergies. Nucleic Acids Res 2020; 48 (W1): W488–w493.

42 Bliss CI. The toxicity of poisons applied jointly 1. Annals of applied biology 1939; 26 (3): 585–615.

43. Webb JL. Enzyme and Metabolic Inhibitors: Academic Press 1963.

44 Fogh J. Cultivation, characterization, and identification of human tumor cells with emphasis on kidney, testis, and bladder tumors. Natl Cancer Inst Monogr 1978 (49): 5–9.

45 Kim ES. Tivozanib: First Global Approval. Drugs 2017; 77 (17): 1917–1923.

46 Grüllich C. Cabozantinib: Multi-kinase Inhibitor of MET, AXL, RET, and VEGFR2. Recent Results Cancer Res 2018; 211: 67-75.

47 Zschäbitz S, Grüllich C. Lenvantinib: A Tyrosine Kinase Inhibitor of VEGFR 1-3, FGFR 1–4, PDGFRα, KIT and RET. Recent Results Cancer Res 2018; 211: 187-198.

48 Rothman J. The Rediscovery of Bisantrene: A Review of the Literature. International Journal of Cancer Research & Therapy 2017; 2 (2): 1–10.

49 Powles T, Albiges L, Bex A et al. Renal cell carcinoma: ESMO Clinical Practice Guideline for diagnosis, treatment and follow-up. Annals of Oncology 2024; 35 (8): 692–706.

50 Acharya N, Singh KP. Differential sensitivity of renal carcinoma cells to doxorubicin and epigenetic therapeutics depends on the genetic background. Mol Cell Biochem 2021; 476 (6): 2365–2379.

51 Gao YH, Wu ZX, Xie LQ et al. VHL deficiency augments anthracycline sensitivity of clear cell renal cell carcinomas by down-regulating ALDH2. Nat Commun 2017; 8: 15337.

52 Gossage L, Eisen T, Maher ER. VHL, the story of a tumour suppressor gene. Nature Reviews Cancer 2015; 15 (1): 55–64.

53 Hwang IY, Roe JS, Seol JH et al. pVHL-mediated transcriptional repression of c-Myc by recruitment of histone deacetylases. Mol Cells 2012; 33 (2): 195–201.

