## Supplementary Data for "Bisantrene potentiates tyrosine kinase inhibitor activity in clear cell renal cell carcinoma"

**Supplementary Data 1**

**Bisantrene potentiates tyrosine kinase inhibitor activity in clear cell renal cell carcinoma**

**Joshua S. Brzozowski^1^, Sumit Sahni^2^, Heather C. Murray^1^, Lauren Watt^1^, Dylan Kiltschewskij^1^, Murray J. Cairns^1^, Marinella Messina^2^, Daniel Tillett^2^, Michael J. Kelso^2^ and Nicole M. Verrills^1^.**

^1^School of Biomedical Sciences and Pharmacy, College of Health, Medicine and Wellbeing, University of Newcastle, Newcastle, New South Wales, Australia and Precision Medicine Program, Hunter Medical Research Institute, New Lambton, New South Wales, Australia.

^2^Race Oncology Limited, Sydney, New South Wales, Australia

Supplementary Table 1: Media composition used for different RCC cell lines.

| **Cell line** | **Base Media** | **Supplements** |
| --- | --- | --- |
| **ACHN, A-704, A-498** | Minimum Essential Medium (MEM) | 1 x non‐essential amino acid solution, 10% foetal bovine serum (FBS), 2 mM L‐glutamine, 1 mM sodium pyruvate |
| **Caki-1, Caki-2** | McCoy's 5A | 10% FBS, 2 mM L‐glutamine |
| **769-P, 786-O** | RPMI‐1640 (with GlutaMAX) | 10% FBS, 20 mM HEPES, 1 mM sodium pyruvate |
| **KMRC-1** | DMEM (high glucose) | 10% FBS, 2 mM L‐glutamine, 20 mM HEPES |
| **RCC4-EV, RCC4-VHL** | DMEM (high glucose) | 10% FBS, 2 mM L‐glutamine, 20 mM HEPES, 0.5mg/mL G418 |

**Supplementary Table 2: Bliss scores for each individual concentration combinations in: (A) 786-O; (B) Caki-1; (C) Caki-2; (D) RCC4-EV; and (E) RCC4-VHL cells.** Values >10 are synergistic (red); values below -10 are antagonistic. Values between-10 to 10 are additive.

| **(A) 786-O** | | | | | | | | | | | | | | | | | | | | | | | | |
| --- | --- | --- | --- | --- | --- | --- | --- | --- | --- | --- | --- | --- | --- | --- | --- | --- | --- | --- | --- | --- | --- | --- | --- | --- |
|  | **Bis:Len** | | | **Bis:Paz** | | | **Bis:Cabo** | | | **Bis:Suni** | | | **Bis:Soraf** | | | **Bis:Axit** | | | **Bis:Tem** | | | **Bis:Ever** | | |
| Bis (µM) | Len  (µM) | RI | SS | Paz  (µM) | RI | SS | Cabo (µM) | RI | SS | Suni (µM) | RI | SS | Soraf (µM) | RI | SS | Axit (µM) | RI | SS | Tem (µM) | RI | SS | Ever (µM) | RI | SS |
| 0 | 0 | 0.000 | 0.000 | 0 | 0.000 | 0.000 | 0 | 0.000 | 0.000 | 0 | 0.000 | 0.000 | 0 | 0.000 | 0.000 | 0 | 0.000 | 0.000 | 0 | 0.000 | 0.000 | 0 | 0.000 | 0.000 |
| 0.5 | 0 | 31.980 | 0.000 | 0 | 37.695 | 0.000 | 0 | 29.519 | 0.000 | 0 | 33.996 | 0.000 | 0 | 33.609 | 0.000 | 0 | 27.605 | 0.000 | 0 | 35.112 | 0.000 | 0 | 34.577 | 0.000 |
| 1 | 0 | 51.291 | 0.000 | 0 | 57.567 | 0.000 | 0 | 49.854 | 0.000 | 0 | 55.089 | 0.000 | 0 | 52.930 | 0.000 | 0 | 47.084 | 0.000 | 0 | 55.727 | 0.000 | 0 | 55.090 | 0.000 |
| 2 | 0 | 72.311 | 0.000 | 0 | 75.099 | 0.000 | 0 | 70.454 | 0.000 | 0 | 71.812 | 0.000 | 0 | 72.191 | 0.000 | 0 | 68.594 | 0.000 | 0 | 72.664 | 0.000 | 0 | 72.759 | 0.000 |
| 4 | 0 | 82.121 | 0.000 | 0 | 81.862 | 0.000 | 0 | 80.856 | 0.000 | 0 | 81.088 | 0.000 | 0 | 81.219 | 0.000 | 0 | 79.906 | 0.000 | 0 | 80.204 | 0.000 | 0 | 81.147 | 0.000 |
| 8 | 0 | 87.474 | 0.000 | 0 | 87.930 | 0.000 | 0 | 86.642 | 0.000 | 0 | 87.047 | 0.000 | 0 | 86.908 | 0.000 | 0 | 86.618 | 0.000 | 0 | 87.150 | 0.000 | 0 | 86.916 | 0.000 |
| 0 | 1.25 | 32.233 | 0.000 | 1.25 | 15.683 | 0.000 | 1.25 | 18.211 | 0.000 | 0.5 | 2.456 | 0.000 | 1.25 | 8.874 | 0.000 | 0.5 | 30.796 | 0.000 | 0.125 | 49.296 | 0.000 | 1.25 | 47.115 | 0.000 |
| 0.5 | 1.25 | 66.432 | 33.097 | 1.25 | 61.555 | 35.575 | 1.25 | 54.401 | 31.918 | 0.5 | 47.982 | 28.796 | 1.25 | 50.063 | 30.220 | 0.5 | 49.126 | 18.180 | 0.125 | 65.748 | 17.115 | 1.25 | 73.418 | 30.884 |
| 1 | 1.25 | 83.290 | 33.127 | 1.25 | 76.676 | 27.848 | 1.25 | 69.524 | 25.143 | 0.5 | 68.415 | 24.109 | 1.25 | 68.117 | 25.761 | 0.5 | 59.171 | 8.568 | 0.125 | 72.398 | 5.711 | 1.25 | 79.293 | 17.747 |
| 2 | 1.25 | 90.095 | 18.309 | 1.25 | 84.737 | 14.347 | 1.25 | 81.230 | 13.807 | 0.5 | 80.314 | 15.295 | 1.25 | 80.695 | 14.643 | 0.5 | 74.802 | 3.835 | 0.125 | 79.633 | -1.162 | 1.25 | 83.231 | 4.855 |
| 4 | 1.25 | 92.900 | 10.917 | 1.25 | 87.988 | 9.318 | 1.25 | 85.526 | 6.083 | 0.5 | 85.922 | 9.230 | 1.25 | 87.296 | 10.301 | 0.5 | 83.069 | 1.405 | 0.125 | 83.721 | -2.999 | 1.25 | 90.387 | 6.158 |
| 8 | 1.25 | 96.208 | 9.169 | 1.25 | 92.010 | 6.205 | 1.25 | 90.097 | 4.488 | 0.5 | 89.160 | 4.902 | 1.25 | 90.176 | 5.989 | 0.5 | 87.305 | -0.903 | 0.125 | 90.875 | 0.087 | 1.25 | 93.661 | 4.741 |
| 0 | 2.5 | 35.174 | 0.000 | 2.5 | 27.243 | 0.000 | 2.5 | 20.662 | 0.000 | 1 | 4.886 | 0.000 | 2.5 | 17.190 | 0.000 | 1 | 38.386 | 0.000 | 0.25 | 48.815 | 0.000 | 2.5 | 47.204 | 0.000 |
| 0.5 | 2.5 | 73.644 | 39.065 | 2.5 | 75.692 | 41.925 | 2.5 | 66.764 | 44.577 | 1 | 52.470 | 31.863 | 2.5 | 55.130 | 28.089 | 1 | 52.493 | 13.319 | 0.25 | 68.081 | 21.026 | 2.5 | 76.398 | 34.953 |
| 1 | 2.5 | 85.762 | 33.937 | 2.5 | 84.153 | 29.450 | 2.5 | 78.749 | 34.681 | 1 | 73.196 | 28.242 | 2.5 | 72.315 | 24.996 | 1 | 64.480 | 8.705 | 0.25 | 75.328 | 10.266 | 2.5 | 81.883 | 21.305 |
| 2 | 2.5 | 91.399 | 18.639 | 2.5 | 86.914 | 12.488 | 2.5 | 84.532 | 16.786 | 1 | 82.948 | 17.456 | 2.5 | 83.407 | 14.477 | 1 | 78.309 | 4.377 | 0.25 | 81.104 | 1.174 | 2.5 | 85.854 | 8.489 |
| 4 | 2.5 | 93.510 | 10.831 | 2.5 | 91.532 | 10.443 | 2.5 | 87.070 | 7.276 | 1 | 86.539 | 9.316 | 2.5 | 89.140 | 10.203 | 1 | 84.680 | 0.932 | 0.25 | 85.122 | -0.833 | 2.5 | 93.135 | 9.983 |
| 8 | 2.5 | 96.644 | 9.120 | 2.5 | 94.204 | 6.745 | 2.5 | 91.194 | 5.344 | 1 | 90.161 | 5.645 | 2.5 | 91.723 | 6.244 | 1 | 88.209 | -1.436 | 0.25 | 92.102 | 1.940 | 2.5 | 93.608 | 4.644 |
| 0 | 5 | 36.467 | 0.000 | 5 | 35.861 | 0.000 | 5 | 28.831 | 0.000 | 2 | 13.347 | 0.000 | 5 | 22.403 | 0.000 | 2 | 45.920 | 0.000 | 0.5 | 48.928 | 0.000 | 5 | 45.804 | 0.000 |
| 0.5 | 5 | 78.285 | 43.578 | 5 | 85.231 | 45.393 | 5 | 80.023 | 52.167 | 2 | 59.985 | 32.923 | 5 | 62.461 | 31.869 | 2 | 60.060 | 13.969 | 0.5 | 70.135 | 23.775 | 5 | 78.357 | 39.509 |
| 1 | 5 | 86.852 | 34.296 | 5 | 87.786 | 28.186 | 5 | 84.620 | 35.671 | 2 | 75.977 | 26.199 | 5 | 76.916 | 26.942 | 2 | 69.410 | 8.401 | 0.5 | 77.184 | 12.782 | 5 | 83.637 | 25.007 |
| 2 | 5 | 92.733 | 19.762 | 5 | 90.301 | 13.335 | 5 | 86.136 | 15.071 | 2 | 83.210 | 14.415 | 5 | 86.574 | 16.179 | 2 | 80.593 | 3.365 | 0.5 | 81.872 | 2.196 | 5 | 90.569 | 15.862 |
| 4 | 5 | 94.008 | 11.089 | 5 | 94.529 | 11.731 | 5 | 90.457 | 9.072 | 2 | 87.157 | 7.801 | 5 | 90.761 | 10.715 | 2 | 86.070 | 0.192 | 0.5 | 87.240 | 2.108 | 5 | 93.600 | 11.156 |
| 8 | 5 | 96.674 | 8.890 | 5 | 96.825 | 8.396 | 5 | 93.939 | 7.071 | 2 | 91.760 | 6.001 | 5 | 93.585 | 7.501 | 2 | 90.431 | -0.250 | 0.5 | 93.233 | 3.506 | 5 | 94.308 | 5.987 |
| 0 | 10 | 38.406 | 0.000 | 10 | 37.287 | 0.000 | 10 | 45.869 | 0.000 | 4 | 56.539 | 0.000 | 10 | 75.074 | 0.000 | 4 | 59.789 | 0.000 | 1 | 50.578 | 0.000 | 10 | 45.009 | 0.000 |
| 0.5 | 10 | 81.491 | 45.520 | 10 | 85.493 | 44.300 | 10 | 85.022 | 39.945 | 4 | 75.120 | 10.812 | 10 | 82.989 | 4.140 | 4 | 66.232 | 5.113 | 1 | 71.552 | 23.646 | 10 | 79.115 | 41.598 |
| 1 | 10 | 87.754 | 33.893 | 10 | 87.967 | 27.444 | 10 | 84.746 | 22.727 | 4 | 82.945 | 7.207 | 10 | 86.578 | 1.259 | 4 | 72.746 | 0.405 | 1 | 78.826 | 13.645 | 10 | 83.051 | 24.888 |
| 2 | 10 | 92.992 | 19.207 | 10 | 92.680 | 15.778 | 10 | 87.878 | 9.511 | 4 | 87.505 | 2.402 | 10 | 89.429 | -2.507 | 4 | 82.830 | -1.048 | 1 | 83.110 | 3.045 | 10 | 93.588 | 20.528 |
| 4 | 10 | 94.926 | 11.697 | 10 | 95.460 | 12.495 | 10 | 91.094 | 4.861 | 4 | 94.079 | 4.529 | 10 | 93.295 | -1.159 | 4 | 87.237 | -2.973 | 1 | 90.325 | 5.809 | 10 | 93.263 | 10.978 |
| 8 | 10 | 96.890 | 8.767 | 10 | 97.437 | 8.896 | 10 | 95.489 | 5.500 | 4 | 96.272 | 3.489 | 10 | 94.890 | -1.345 | 4 | 91.863 | -1.510 | 1 | 94.011 | 4.181 | 10 | 95.207 | 7.455 |
| 0 | 20 | 25.360 | 0.000 | 20 | 15.729 | 0.000 | 20 | 63.799 | 0.000 | 8 | 91.472 | 0.000 | 20 | 88.445 | 0.000 | 8 | 53.861 | 0.000 | 2 | 46.958 | 0.000 | 20 | 49.221 | 0.000 |
| 0.5 | 20 | 77.554 | 55.141 | 20 | 71.580 | 48.223 | 20 | 86.290 | 22.141 | 8 | 92.825 | -0.596 | 20 | 87.721 | -3.490 | 8 | 64.088 | 9.536 | 2 | 72.982 | 30.331 | 20 | 77.441 | 33.816 |
| 1 | 20 | 84.733 | 40.524 | 20 | 81.368 | 33.760 | 20 | 85.282 | 9.607 | 8 | 96.086 | 0.742 | 20 | 89.082 | -5.198 | 8 | 74.714 | 8.219 | 2 | 78.253 | 16.024 | 20 | 88.106 | 28.257 |
| 2 | 20 | 91.476 | 23.234 | 20 | 87.270 | 17.537 | 20 | 92.532 | 7.151 | 8 | 98.923 | 2.101 | 20 | 92.800 | -3.994 | 8 | 82.095 | 1.125 | 2 | 84.817 | 7.419 | 20 | 91.013 | 14.651 |
| 4 | 20 | 95.261 | 15.988 | 20 | 92.391 | 14.880 | 20 | 93.951 | 3.135 | 8 | 99.299 | 1.434 | 20 | 95.587 | -2.143 | 8 | 88.284 | 0.382 | 2 | 93.328 | 11.468 | 20 | 95.203 | 12.137 |
| 8 | 20 | 97.473 | 12.221 | 20 | 95.826 | 11.029 | 20 | 97.283 | 4.050 | 8 | 99.428 | 0.874 | 20 | 97.644 | -0.608 | 8 | 93.582 | 2.049 | 2 | 95.076 | 6.607 | 20 | 98.034 | 10.337 |
| **(B) Caki-1** | | | | | | | | | | | | | | | | | | | | | | | | |
|  | **Bis:Len** | | | **Bis:Paz** | | | **Bis:Cabo** | | | **Bis:Suni** | | | **Bis:Soraf** | | | **Bis:Axit** | | | **Bis:Tem** | | | **Bis:Ever** | | |
| Bis (µM) | Len  (µM) | RI | SS | Paz  (µM) | RI | SS | Cabo (µM) | RI | SS | Suni (µM) | RI | SS | Soraf (µM) | RI | SS | Axit (µM) | RI | SS | Tem (µM) | RI | SS | Ever (µM) | RI | SS |
| 0 | 0 | 0.000 | 0.000 | 0 | 0.000 | 0.000 | 0 | 0.000 | 0.000 | 0 | 0.000 | 0.000 | 0 | 0.000 | 0.000 | 0 | 0.000 | 0.000 | 0 | 0.000 | 0.000 | 0 | 0.000 | 0.000 |
| 0.25 | 0 | 40.497 | 0.000 | 0 | 46.192 | 0.000 | 0 | 43.480 | 0.000 | 0 | 32.961 | 0.000 | 0 | 32.161 | 0.000 | 0 | 34.687 | 0.000 | 0 | 28.721 | 0.000 | 0 | 29.029 | 0.000 |
| 0.5 | 0 | 56.467 | 0.000 | 0 | 61.696 | 0.000 | 0 | 57.989 | 0.000 | 0 | 46.755 | 0.000 | 0 | 45.895 | 0.000 | 0 | 50.558 | 0.000 | 0 | 42.081 | 0.000 | 0 | 43.032 | 0.000 |
| 1 | 0 | 70.238 | 0.000 | 0 | 74.729 | 0.000 | 0 | 70.140 | 0.000 | 0 | 60.838 | 0.000 | 0 | 60.400 | 0.000 | 0 | 66.607 | 0.000 | 0 | 58.864 | 0.000 | 0 | 58.392 | 0.000 |
| 2 | 0 | 80.092 | 0.000 | 0 | 83.222 | 0.000 | 0 | 80.753 | 0.000 | 0 | 73.910 | 0.000 | 0 | 72.709 | 0.000 | 0 | 78.281 | 0.000 | 0 | 71.779 | 0.000 | 0 | 71.474 | 0.000 |
| 4 | 0 | 80.743 | 0.000 | 0 | 83.980 | 0.000 | 0 | 80.357 | 0.000 | 0 | 80.119 | 0.000 | 0 | 80.881 | 0.000 | 0 | 77.422 | 0.000 | 0 | 78.979 | 0.000 | 0 | 77.900 | 0.000 |
| 0 | 1.25 | 9.875 | 0.000 | 1.25 | 13.107 | 0.000 | 1.25 | 6.949 | 0.000 | 0.5 | 10.040 | 0.000 | 1.25 | 6.790 | 0.000 | 0.5 | 4.542 | 0.000 | 0.125 | 32.348 | 0.000 | 1.25 | 33.573 | 0.000 |
| 0.25 | 1.25 | 59.758 | 33.553 | 1.25 | 64.412 | 32.475 | 1.25 | 58.823 | 31.211 | 0.5 | 43.271 | 20.110 | 1.25 | 43.247 | 23.486 | 0.5 | 42.449 | 20.415 | 0.125 | 54.951 | 23.358 | 1.25 | 61.725 | 29.541 |
| 0.5 | 1.25 | 74.164 | 29.057 | 1.25 | 75.104 | 23.686 | 1.25 | 70.943 | 25.145 | 0.5 | 54.601 | 15.556 | 1.25 | 57.876 | 22.516 | 0.5 | 57.478 | 16.704 | 0.125 | 63.837 | 19.558 | 1.25 | 67.445 | 21.338 |
| 1 | 1.25 | 84.950 | 23.134 | 1.25 | 82.362 | 14.053 | 1.25 | 78.366 | 16.642 | 0.5 | 65.114 | 9.626 | 1.25 | 68.610 | 15.806 | 0.5 | 71.289 | 11.291 | 0.125 | 71.821 | 10.634 | 1.25 | 76.077 | 15.397 |
| 2 | 1.25 | 85.574 | 10.023 | 1.25 | 87.644 | 8.493 | 1.25 | 86.054 | 10.723 | 0.5 | 75.165 | 4.477 | 1.25 | 77.178 | 9.459 | 0.5 | 81.794 | 7.904 | 0.125 | 79.795 | 6.156 | 1.25 | 83.379 | 10.272 |
| 4 | 1.25 | 85.601 | 9.139 | 1.25 | 87.691 | 7.446 | 1.25 | 86.015 | 11.256 | 0.5 | 80.037 | 2.150 | 1.25 | 80.777 | 2.727 | 0.5 | 82.263 | 9.638 | 0.125 | 83.022 | 2.069 | 1.25 | 84.199 | 4.170 |
| 0 | 2.5 | 10.640 | 0.000 | 2.5 | 26.470 | 0.000 | 2.5 | 12.661 | 0.000 | 1 | 17.160 | 0.000 | 2.5 | 12.611 | 0.000 | 1 | 11.334 | 0.000 | 0.25 | 34.358 | 0.000 | 2.5 | 34.624 | 0.000 |
| 0.25 | 2.5 | 63.591 | 37.636 | 2.5 | 74.754 | 33.794 | 2.5 | 67.302 | 36.768 | 1 | 50.117 | 21.381 | 2.5 | 48.284 | 23.819 | 1 | 46.876 | 19.359 | 0.25 | 57.091 | 23.710 | 2.5 | 63.544 | 30.646 |
| 0.5 | 2.5 | 77.589 | 32.819 | 2.5 | 81.969 | 23.980 | 2.5 | 76.035 | 27.755 | 1 | 59.433 | 15.830 | 2.5 | 60.788 | 21.449 | 1 | 61.852 | 17.128 | 0.25 | 65.537 | 19.793 | 2.5 | 69.711 | 23.270 |
| 1 | 2.5 | 88.461 | 27.170 | 2.5 | 86.899 | 14.257 | 2.5 | 82.538 | 19.191 | 1 | 69.462 | 10.792 | 2.5 | 68.973 | 12.893 | 1 | 74.405 | 11.771 | 0.25 | 72.936 | 10.680 | 2.5 | 77.666 | 16.722 |
| 2 | 2.5 | 87.840 | 12.618 | 2.5 | 89.159 | 6.687 | 2.5 | 88.014 | 11.451 | 1 | 78.192 | 5.411 | 2.5 | 78.151 | 8.324 | 1 | 83.883 | 8.291 | 0.25 | 81.490 | 7.401 | 2.5 | 85.714 | 12.794 |
| 4 | 2.5 | 87.334 | 11.076 | 2.5 | 88.226 | 4.540 | 2.5 | 87.315 | 11.120 | 1 | 80.268 | 0.340 | 2.5 | 81.896 | 2.458 | 1 | 84.268 | 9.842 | 0.25 | 84.126 | 2.791 | 2.5 | 84.484 | 4.149 |
| 0 | 5 | 18.539 | 0.000 | 5 | 42.708 | 0.000 | 5 | 23.703 | 0.000 | 2 | 28.379 | 0.000 | 5 | 17.141 | 0.000 | 2 | 17.887 | 0.000 | 0.5 | 32.688 | 0.000 | 5 | 33.198 | 0.000 |
| 0.25 | 5 | 67.701 | 35.423 | 5 | 82.175 | 28.725 | 5 | 75.371 | 37.061 | 2 | 54.374 | 15.450 | 5 | 48.797 | 19.970 | 2 | 48.344 | 14.987 | 0.5 | 60.215 | 29.816 | 5 | 66.838 | 36.613 |
| 0.5 | 5 | 78.725 | 28.862 | 5 | 86.246 | 19.068 | 5 | 81.008 | 26.687 | 2 | 63.210 | 11.600 | 5 | 58.915 | 15.628 | 2 | 63.922 | 14.966 | 0.5 | 67.167 | 23.569 | 5 | 72.645 | 28.435 |
| 1 | 5 | 89.544 | 24.846 | 5 | 90.812 | 12.431 | 5 | 86.894 | 19.463 | 2 | 72.360 | 7.826 | 5 | 70.702 | 12.370 | 2 | 76.388 | 11.011 | 0.5 | 75.198 | 14.804 | 5 | 81.255 | 22.367 |
| 2 | 5 | 88.100 | 10.484 | 5 | 90.860 | 4.312 | 5 | 89.016 | 9.359 | 2 | 79.748 | 2.981 | 5 | 78.699 | 7.187 | 2 | 85.047 | 7.646 | 0.5 | 82.721 | 9.811 | 5 | 86.903 | 15.017 |
| 4 | 5 | 88.871 | 10.618 | 5 | 90.281 | 2.831 | 5 | 88.924 | 9.720 | 2 | 82.311 | -0.471 | 5 | 84.167 | 3.938 | 2 | 84.760 | 8.312 | 0.5 | 84.595 | 4.001 | 5 | 84.271 | 4.402 |
| 0 | 10 | 25.339 | 0.000 | 10 | 52.030 | 0.000 | 10 | 49.254 | 0.000 | 4 | 38.714 | 0.000 | 10 | 52.600 | 0.000 | 4 | 21.273 | 0.000 | 1 | 34.633 | 0.000 | 10 | 32.484 | 0.000 |
| 0.25 | 10 | 69.277 | 31.066 | 10 | 86.452 | 25.836 | 10 | 80.452 | 20.673 | 4 | 57.105 | 8.539 | 10 | 66.453 | 6.293 | 4 | 48.484 | 11.989 | 1 | 62.734 | 30.743 | 10 | 64.244 | 34.103 |
| 0.5 | 10 | 78.966 | 24.533 | 10 | 88.907 | 16.513 | 10 | 85.332 | 15.193 | 4 | 64.119 | 4.579 | 10 | 71.373 | 2.763 | 4 | 62.753 | 11.169 | 1 | 68.226 | 23.032 | 10 | 71.751 | 27.959 |
| 1 | 10 | 90.746 | 23.184 | 10 | 93.538 | 12.005 | 10 | 91.188 | 12.825 | 4 | 73.541 | 3.285 | 10 | 76.161 | -1.458 | 4 | 75.617 | 8.469 | 1 | 77.178 | 16.026 | 10 | 80.511 | 21.902 |
| 2 | 10 | 87.454 | 7.558 | 10 | 91.692 | 2.761 | 10 | 87.870 | 0.168 | 4 | 80.234 | -0.410 | 10 | 82.653 | -2.111 | 4 | 84.145 | 5.513 | 1 | 83.996 | 10.541 | 10 | 85.972 | 14.153 |
| 4 | 10 | 89.225 | 9.013 | 10 | 92.903 | 3.720 | 10 | 90.306 | 3.543 | 4 | 83.885 | -1.592 | 10 | 85.292 | -4.560 | 4 | 84.000 | 6.308 | 1 | 84.904 | 3.708 | 10 | 84.370 | 4.795 |
| 0 | 20 | 27.805 | 0.000 | 20 | 21.000 | 0.000 | 20 | 72.170 | 0.000 | 8 | 46.990 | 0.000 | 20 | 83.460 | 0.000 | 8 | 25.021 | 0.000 | 2 | 32.957 | 0.000 | 20 | 30.004 | 0.000 |
| 0.25 | 20 | 65.836 | 24.467 | 20 | 75.480 | 39.685 | 20 | 85.573 | 6.683 | 8 | 56.347 | -0.573 | 20 | 85.668 | -0.969 | 8 | 48.217 | 8.162 | 2 | 60.949 | 30.448 | 20 | 62.772 | 35.149 |
| 0.5 | 20 | 74.094 | 16.760 | 20 | 81.056 | 26.320 | 20 | 88.397 | 3.864 | 8 | 61.724 | -4.836 | 20 | 86.993 | -2.674 | 8 | 57.308 | 2.000 | 2 | 67.330 | 23.516 | 20 | 70.369 | 28.540 |
| 1 | 20 | 88.468 | 19.187 | 20 | 88.481 | 18.635 | 20 | 92.267 | 3.391 | 8 | 73.998 | -0.944 | 20 | 87.728 | -5.275 | 8 | 75.892 | 7.006 | 2 | 77.328 | 17.396 | 20 | 80.946 | 24.198 |
| 2 | 20 | 85.391 | 4.210 | 20 | 88.577 | 7.476 | 20 | 86.883 | -8.026 | 8 | 85.647 | 2.979 | 20 | 90.080 | -5.399 | 8 | 82.540 | 2.427 | 2 | 85.235 | 12.963 | 20 | 82.834 | 11.275 |
| 4 | 20 | 88.254 | 7.056 | 20 | 89.521 | 7.693 | 20 | 90.544 | -3.254 | 8 | 93.027 | 7.087 | 20 | 90.520 | -6.833 | 8 | 82.603 | 3.424 | 2 | 84.695 | 4.034 | 20 | 84.885 | 6.382 |
| **(C) Caki-2** | | | | | | | | | | | | | | | | | | | | | | | | |
|  | **Bis:Len** | | | **Bis:Paz** | | | **Bis:Cabo** | | | **Bis:Suni** | | | **Bis:Soraf** | | | **Bis:Axit** | | | **Bis:Tem** | | | **Bis:Ever** | | |
| Bis (µM) | Len  (µM) | RI | SS | Paz  (µM) | RI | SS | Cabo (µM) | RI | SS | Suni (µM) | RI | SS | Soraf (µM) | RI | SS | Axit (µM) | RI | SS | Tem (µM) | RI | SS | Ever (µM) | RI | SS |
| 0 | 0 | 0.000 | 0.000 | 0 | 0.000 | 0.000 | 0 | 0.000 | 0.000 | 0 | 0.000 | 0.000 | 0 | 0.000 | 0.000 | 0 | 0.000 | 0.000 | 0 | 0.000 | 0.000 | 0 | 0.000 | 0.000 |
| 0.125 | 0 | 22.920 | 0.000 | 0 | 27.891 | 0.000 | 0 | 29.316 | 0.000 | 0 | 28.226 | 0.000 | 0 | 29.132 | 0.000 | 0 | 21.350 | 0.000 | 0 | 28.466 | 0.000 | 0 | 26.907 | 0.000 |
| 0.25 | 0 | 37.620 | 0.000 | 0 | 45.181 | 0.000 | 0 | 41.449 | 0.000 | 0 | 46.898 | 0.000 | 0 | 47.430 | 0.000 | 0 | 34.012 | 0.000 | 0 | 45.412 | 0.000 | 0 | 46.024 | 0.000 |
| 0.5 | 0 | 46.878 | 0.000 | 0 | 56.514 | 0.000 | 0 | 51.893 | 0.000 | 0 | 57.375 | 0.000 | 0 | 57.772 | 0.000 | 0 | 44.188 | 0.000 | 0 | 56.160 | 0.000 | 0 | 55.698 | 0.000 |
| 1 | 0 | 63.915 | 0.000 | 0 | 72.883 | 0.000 | 0 | 68.557 | 0.000 | 0 | 69.529 | 0.000 | 0 | 68.891 | 0.000 | 0 | 62.167 | 0.000 | 0 | 67.561 | 0.000 | 0 | 68.568 | 0.000 |
| 2 | 0 | 73.779 | 0.000 | 0 | 76.320 | 0.000 | 0 | 77.017 | 0.000 | 0 | 79.361 | 0.000 | 0 | 79.381 | 0.000 | 0 | 72.114 | 0.000 | 0 | 79.017 | 0.000 | 0 | 78.829 | 0.000 |
| 0 | 1.25 | 18.923 | 0.000 | 1.25 | 5.929 | 0.000 | 1.25 | 24.803 | 0.000 | 0.5 | 4.192 | 0.000 | 1.25 | -7.697 | 0.000 | 0.5 | 24.965 | 0.000 | 0.125 | 35.619 | 0.000 | 1.25 | 35.646 | 0.000 |
| 0.125 | 1.25 | 47.855 | 28.670 | 1.25 | 32.962 | 15.388 | 1.25 | 50.043 | 22.742 | 0.5 | 35.375 | 18.047 | 1.25 | 24.613 | 9.706 | 0.5 | 36.592 | 9.362 | 0.125 | 55.895 | 22.298 | 1.25 | 58.577 | 27.075 |
| 0.25 | 1.25 | 60.689 | 26.707 | 1.25 | 53.314 | 16.747 | 1.25 | 59.631 | 20.129 | 0.5 | 52.760 | 14.020 | 1.25 | 46.529 | 9.904 | 0.5 | 48.071 | 9.387 | 0.125 | 65.366 | 15.722 | 1.25 | 69.453 | 20.049 |
| 0.5 | 1.25 | 65.510 | 21.514 | 1.25 | 62.866 | 13.167 | 1.25 | 64.991 | 14.191 | 0.5 | 63.937 | 13.417 | 1.25 | 60.198 | 11.429 | 0.5 | 52.066 | 3.101 | 0.125 | 69.875 | 9.569 | 1.25 | 72.755 | 13.593 |
| 1 | 1.25 | 81.757 | 20.903 | 1.25 | 79.974 | 11.905 | 1.25 | 74.441 | 5.862 | 0.5 | 73.038 | 8.215 | 1.25 | 69.886 | 7.546 | 0.5 | 66.312 | 0.660 | 0.125 | 73.749 | 1.840 | 1.25 | 78.221 | 6.418 |
| 2 | 1.25 | 79.808 | 6.778 | 1.25 | 82.652 | 10.561 | 1.25 | 79.294 | 1.703 | 0.5 | 77.176 | 0.249 | 1.25 | 80.751 | 5.786 | 0.5 | 74.228 | -0.649 | 0.125 | 79.422 | -3.572 | 1.25 | 80.353 | -2.236 |
| 0 | 2.5 | 34.182 | 0.000 | 2.5 | 9.796 | 0.000 | 2.5 | 37.527 | 0.000 | 1 | 13.105 | 0.000 | 2.5 | -8.402 | 0.000 | 1 | 39.374 | 0.000 | 0.25 | 35.374 | 0.000 | 2.5 | 35.741 | 0.000 |
| 0.125 | 2.5 | 57.442 | 22.999 | 2.5 | 39.111 | 18.760 | 2.5 | 61.174 | 22.308 | 1 | 45.929 | 21.660 | 2.5 | 26.382 | 12.267 | 1 | 45.344 | 3.435 | 0.25 | 56.620 | 23.563 | 2.5 | 62.047 | 31.526 |
| 0.25 | 2.5 | 68.246 | 21.872 | 2.5 | 59.980 | 21.662 | 2.5 | 68.369 | 19.156 | 1 | 62.653 | 19.122 | 2.5 | 49.090 | 13.180 | 1 | 53.257 | 1.813 | 0.25 | 67.975 | 19.408 | 2.5 | 72.625 | 24.139 |
| 0.5 | 2.5 | 71.121 | 16.397 | 2.5 | 66.135 | 14.683 | 2.5 | 72.221 | 13.455 | 1 | 69.878 | 15.182 | 2.5 | 61.667 | 13.412 | 1 | 58.143 | -1.268 | 0.25 | 71.131 | 11.419 | 2.5 | 75.386 | 16.986 |
| 1 | 2.5 | 84.035 | 15.568 | 2.5 | 83.006 | 14.023 | 2.5 | 78.265 | 4.232 | 1 | 73.528 | 5.100 | 2.5 | 70.772 | 8.792 | 1 | 68.592 | -4.518 | 0.25 | 75.245 | 3.959 | 2.5 | 80.902 | 9.898 |
| 2 | 2.5 | 79.928 | 1.039 | 2.5 | 85.625 | 12.795 | 2.5 | 80.494 | -1.523 | 1 | 77.271 | -2.137 | 2.5 | 81.269 | 6.534 | 1 | 74.248 | -6.468 | 0.25 | 80.017 | -2.694 | 2.5 | 81.030 | -1.379 |
| 0 | 5 | 38.751 | 0.000 | 5 | 19.584 | 0.000 | 5 | 46.556 | 0.000 | 2 | 29.235 | 0.000 | 5 | -11.859 | 0.000 | 2 | 42.425 | 0.000 | 0.5 | 35.918 | 0.000 | 5 | 36.043 | 0.000 |
| 0.125 | 5 | 60.007 | 20.930 | 5 | 48.876 | 20.453 | 5 | 69.234 | 22.207 | 2 | 55.074 | 16.600 | 5 | 23.176 | 11.710 | 2 | 50.028 | 5.594 | 0.5 | 57.270 | 23.743 | 5 | 62.945 | 32.326 |
| 0.25 | 5 | 69.617 | 19.343 | 5 | 63.723 | 18.605 | 5 | 75.899 | 20.161 | 2 | 67.679 | 13.351 | 5 | 44.581 | 10.417 | 2 | 56.427 | 2.708 | 0.5 | 67.859 | 18.735 | 5 | 72.489 | 23.677 |
| 0.5 | 5 | 71.040 | 12.729 | 5 | 69.435 | 12.648 | 5 | 77.007 | 12.495 | 2 | 73.630 | 10.221 | 5 | 58.614 | 11.820 | 2 | 58.516 | -3.293 | 0.5 | 72.628 | 12.983 | 5 | 76.067 | 17.651 |
| 1 | 5 | 86.894 | 16.610 | 5 | 86.545 | 14.503 | 5 | 82.461 | 4.965 | 2 | 76.635 | 2.048 | 5 | 70.610 | 9.921 | 2 | 69.828 | -4.706 | 0.5 | 76.996 | 5.967 | 5 | 84.177 | 14.048 |
| 2 | 5 | 79.517 | -1.221 | 5 | 88.541 | 13.008 | 5 | 84.391 | 0.073 | 2 | 78.461 | -5.269 | 5 | 82.961 | 9.268 | 2 | 74.801 | -7.039 | 0.5 | 80.516 | -2.234 | 5 | 81.172 | -1.303 |
| 0 | 10 | 32.139 | 0.000 | 10 | 39.757 | 0.000 | 10 | 63.394 | 0.000 | 4 | 44.837 | 0.000 | 10 | 13.439 | 0.000 | 4 | 32.991 | 0.000 | 1 | 37.891 | 0.000 | 10 | 38.065 | 0.000 |
| 0.125 | 10 | 58.552 | 26.661 | 10 | 58.901 | 12.016 | 10 | 76.674 | 12.337 | 4 | 59.144 | 6.142 | 10 | 31.568 | -0.846 | 4 | 41.142 | 5.669 | 1 | 58.246 | 22.563 | 10 | 61.233 | 27.505 |
| 0.25 | 10 | 66.906 | 22.122 | 10 | 69.629 | 10.167 | 10 | 79.788 | 9.085 | 4 | 65.715 | -0.183 | 10 | 42.142 | -8.453 | 4 | 49.037 | 2.850 | 1 | 66.233 | 14.697 | 10 | 69.537 | 17.893 |
| 0.5 | 10 | 66.826 | 12.785 | 10 | 72.809 | 4.407 | 10 | 80.346 | 3.578 | 4 | 70.177 | -2.829 | 10 | 52.581 | -7.831 | 4 | 51.979 | -3.519 | 1 | 70.742 | 8.972 | 10 | 72.588 | 11.512 |
| 1 | 10 | 87.411 | 20.746 | 10 | 91.693 | 12.957 | 10 | 87.838 | 3.211 | 4 | 75.675 | -5.525 | 10 | 71.491 | 1.317 | 4 | 69.382 | -0.053 | 1 | 76.810 | 4.600 | 10 | 87.145 | 16.855 |
| 2 | 10 | 77.647 | -0.939 | 10 | 89.198 | 7.127 | 10 | 86.551 | -3.468 | 4 | 79.267 | -8.701 | 10 | 85.198 | 5.389 | 4 | 76.324 | -1.376 | 1 | 80.941 | -2.397 | 10 | 79.006 | -4.900 |
| 0 | 20 | 19.173 | 0.000 | 20 | 13.750 | 0.000 | 20 | 77.572 | 0.000 | 8 | 56.364 | 0.000 | 20 | 81.635 | 0.000 | 8 | 19.338 | 0.000 | 2 | 37.677 | 0.000 | 20 | 37.668 | 0.000 |
| 0.125 | 20 | 50.682 | 31.814 | 20 | 43.770 | 20.287 | 20 | 81.363 | 2.017 | 8 | 60.158 | -3.907 | 20 | 79.738 | -6.515 | 8 | 31.008 | 9.065 | 2 | 55.511 | 19.211 | 20 | 60.012 | 26.400 |
| 0.25 | 20 | 58.883 | 24.287 | 20 | 56.387 | 14.400 | 20 | 81.976 | -1.628 | 8 | 63.650 | -10.898 | 20 | 81.810 | -8.318 | 8 | 36.201 | 0.473 | 2 | 62.597 | 10.089 | 20 | 65.177 | 12.520 |
| 0.5 | 20 | 60.196 | 14.874 | 20 | 62.688 | 8.218 | 20 | 82.383 | -4.920 | 8 | 75.573 | -3.212 | 20 | 86.903 | -5.011 | 8 | 43.056 | -3.198 | 2 | 65.978 | 2.832 | 20 | 70.561 | 9.147 |
| 1 | 20 | 86.901 | 27.008 | 20 | 86.146 | 16.235 | 20 | 94.909 | 4.977 | 8 | 89.274 | 5.551 | 20 | 93.930 | 0.256 | 8 | 69.472 | 7.567 | 2 | 78.354 | 6.764 | 20 | 90.331 | 21.269 |
| 2 | 20 | 77.596 | 3.999 | 20 | 85.561 | 11.415 | 20 | 84.723 | -11.099 | 8 | 90.295 | 0.917 | 20 | 93.970 | -2.044 | 8 | 77.153 | 5.161 | 2 | 80.044 | -3.505 | 20 | 75.373 | -9.540 |
| **(D) RCC4-EV** | | | | | | | | | | | | | | | | | | | | | | | | |
|  | **Bis:Len** | | | **Bis:Paz** | | | **Bis:Cabo** | | | **Bis:Suni** | | | **Bis:Soraf** | | | **Bis:Axit** | | | **Bis:Tem** | | | **Bis:Ever** | | |
| Bis (µM) | Len  (µM) | RI | SS | Paz  (µM) | RI | SS | Cabo (µM) | RI | SS | Suni (µM) | RI | SS | Soraf (µM) | RI | SS | Axit (µM) | RI | SS | Tem (µM) | RI | SS | Ever (µM) | RI | SS |
| 0 | 0 | 0.000 | 0.000 | 0 | 0.000 | 0.000 | 0 | 0.000 | 0.000 | 0 | 0.000 | 0.000 | 0 | 0.000 | 0.000 | 0 | 0.000 | 0.000 | 0 | 0.000 | 0.000 | 0 | 0.000 | 0.000 |
| 0.25 | 0 | 23.236 | 0.000 | 0 | 22.169 | 0.000 | 0 | 24.409 | 0.000 | 0 | 11.660 | 0.000 | 0 | 20.305 | 0.000 | 0 | 22.347 | 0.000 | 0 | 18.107 | 0.000 | 0 | 19.321 | 0.000 |
| 0.5 | 0 | 31.073 | 0.000 | 0 | 38.489 | 0.000 | 0 | 36.868 | 0.000 | 0 | 32.348 | 0.000 | 0 | 33.940 | 0.000 | 0 | 32.333 | 0.000 | 0 | 31.243 | 0.000 | 0 | 32.741 | 0.000 |
| 1 | 0 | 57.196 | 0.000 | 0 | 56.359 | 0.000 | 0 | 56.902 | 0.000 | 0 | 55.736 | 0.000 | 0 | 56.601 | 0.000 | 0 | 51.694 | 0.000 | 0 | 54.203 | 0.000 | 0 | 57.454 | 0.000 |
| 2 | 0 | 69.495 | 0.000 | 0 | 65.947 | 0.000 | 0 | 68.672 | 0.000 | 0 | 65.024 | 0.000 | 0 | 64.169 | 0.000 | 0 | 67.555 | 0.000 | 0 | 61.802 | 0.000 | 0 | 63.382 | 0.000 |
| 4 | 0 | 75.967 | 0.000 | 0 | 75.976 | 0.000 | 0 | 76.131 | 0.000 | 0 | 74.515 | 0.000 | 0 | 74.936 | 0.000 | 0 | 74.201 | 0.000 | 0 | 71.918 | 0.000 | 0 | 73.110 | 0.000 |
| 0 | 1.25 | 13.507 | 0.000 | 1.25 | 12.548 | 0.000 | 1.25 | 14.393 | 0.000 | 0.5 | -8.424 | 0.000 | 1.25 | -3.444 | 0.000 | 0.5 | 18.116 | 0.000 | 0.125 | 53.825 | 0.000 | 1.25 | 53.149 | 0.000 |
| 0.25 | 1.25 | 44.677 | 27.419 | 1.25 | 42.007 | 25.682 | 1.25 | 45.772 | 27.536 | 0.5 | 20.945 | 19.788 | 1.25 | 25.297 | 15.477 | 0.5 | 36.870 | 16.012 | 0.125 | 62.048 | 18.028 | 1.25 | 67.467 | 25.687 |
| 0.5 | 1.25 | 68.364 | 45.950 | 1.25 | 60.731 | 27.999 | 1.25 | 55.956 | 24.485 | 0.5 | 42.409 | 18.183 | 1.25 | 41.748 | 16.792 | 0.5 | 50.314 | 20.356 | 0.125 | 68.160 | 15.169 | 1.25 | 72.074 | 20.319 |
| 1 | 1.25 | 85.851 | 35.035 | 1.25 | 72.235 | 19.978 | 1.25 | 74.173 | 21.780 | 0.5 | 59.720 | 9.228 | 1.25 | 61.020 | 10.265 | 0.5 | 60.274 | 9.414 | 0.125 | 73.092 | 2.377 | 1.25 | 74.504 | 2.202 |
| 2 | 1.25 | 95.865 | 32.002 | 1.25 | 75.873 | 12.700 | 1.25 | 79.216 | 13.431 | 0.5 | 64.360 | 3.378 | 1.25 | 68.305 | 9.002 | 0.5 | 73.625 | 6.693 | 0.125 | 74.390 | -2.309 | 1.25 | 78.403 | 2.367 |
| 4 | 1.25 | 97.552 | 26.170 | 1.25 | 88.964 | 15.983 | 1.25 | 87.416 | 14.116 | 0.5 | 76.735 | 5.236 | 1.25 | 78.057 | 6.543 | 0.5 | 77.534 | 3.526 | 0.125 | 82.534 | 0.147 | 1.25 | 90.545 | 10.449 |
| 0 | 2.5 | 16.494 | 0.000 | 2.5 | 24.415 | 0.000 | 2.5 | 24.651 | 0.000 | 1 | 1.894 | 0.000 | 2.5 | 3.839 | 0.000 | 1 | 24.028 | 0.000 | 0.25 | 53.264 | 0.000 | 2.5 | 54.050 | 0.000 |
| 0.25 | 2.5 | 50.639 | 31.263 | 2.5 | 60.307 | 34.438 | 2.5 | 57.465 | 30.442 | 1 | 31.840 | 21.366 | 2.5 | 32.361 | 16.337 | 1 | 44.257 | 18.258 | 0.25 | 64.086 | 21.634 | 2.5 | 68.745 | 26.080 |
| 0.5 | 2.5 | 74.531 | 50.363 | 2.5 | 72.233 | 31.446 | 2.5 | 72.342 | 34.818 | 1 | 50.863 | 19.505 | 2.5 | 46.390 | 16.194 | 1 | 51.831 | 16.397 | 0.25 | 70.161 | 18.590 | 2.5 | 73.872 | 21.644 |
| 1 | 2.5 | 92.540 | 41.158 | 2.5 | 76.439 | 17.779 | 2.5 | 79.242 | 21.527 | 1 | 60.989 | 5.718 | 2.5 | 61.153 | 6.722 | 1 | 63.010 | 8.576 | 0.25 | 74.222 | 4.382 | 2.5 | 77.932 | 6.154 |
| 2 | 2.5 | 96.229 | 31.158 | 2.5 | 84.467 | 17.220 | 2.5 | 82.152 | 12.354 | 1 | 66.155 | 1.418 | 2.5 | 70.791 | 8.642 | 1 | 74.160 | 4.563 | 0.25 | 75.360 | -0.600 | 2.5 | 85.025 | 10.764 |
| 4 | 2.5 | 96.993 | 24.506 | 2.5 | 94.932 | 19.060 | 2.5 | 95.555 | 20.344 | 1 | 82.337 | 8.217 | 2.5 | 83.041 | 9.796 | 1 | 80.399 | 4.766 | 0.25 | 84.273 | 2.794 | 2.5 | 92.029 | 12.019 |
| 0 | 5 | 18.784 | 0.000 | 5 | 41.267 | 0.000 | 5 | 40.359 | 0.000 | 2 | 17.429 | 0.000 | 5 | 22.829 | 0.000 | 2 | 42.307 | 0.000 | 0.5 | 53.141 | 0.000 | 5 | 54.174 | 0.000 |
| 0.25 | 5 | 59.206 | 38.934 | 5 | 74.715 | 33.283 | 5 | 73.709 | 32.904 | 2 | 43.118 | 18.480 | 5 | 36.273 | 2.909 | 2 | 54.415 | 9.952 | 0.5 | 64.281 | 22.084 | 5 | 70.531 | 28.323 |
| 0.5 | 5 | 78.024 | 52.286 | 5 | 80.538 | 26.916 | 5 | 80.954 | 30.957 | 2 | 59.120 | 16.904 | 5 | 50.884 | 6.415 | 2 | 57.896 | 5.809 | 0.5 | 71.880 | 21.075 | 5 | 75.345 | 23.493 |
| 1 | 5 | 95.087 | 42.798 | 5 | 84.138 | 16.686 | 5 | 83.753 | 17.258 | 2 | 66.305 | 3.929 | 5 | 66.199 | 2.558 | 2 | 66.747 | 0.305 | 0.5 | 75.785 | 6.603 | 5 | 79.303 | 7.921 |
| 2 | 5 | 96.075 | 29.999 | 5 | 96.888 | 23.894 | 5 | 91.584 | 16.599 | 2 | 69.740 | -0.628 | 5 | 76.570 | 6.948 | 2 | 74.469 | -3.639 | 0.5 | 77.018 | 1.733 | 5 | 92.228 | 20.478 |
| 4 | 5 | 96.934 | 23.665 | 5 | 97.782 | 16.830 | 5 | 97.232 | 16.999 | 2 | 90.107 | 12.021 | 5 | 89.226 | 10.924 | 2 | 82.415 | 0.372 | 0.5 | 87.122 | 6.719 | 5 | 90.568 | 9.970 |
| 0 | 10 | 23.099 | 0.000 | 10 | 59.048 | 0.000 | 10 | 60.739 | 0.000 | 4 | 38.684 | 0.000 | 10 | 49.159 | 0.000 | 4 | 47.334 | 0.000 | 1 | 53.683 | 0.000 | 10 | 54.134 | 0.000 |
| 0.25 | 10 | 64.827 | 40.950 | 10 | 82.718 | 23.615 | 10 | 81.496 | 20.231 | 4 | 55.288 | 11.185 | 10 | 55.176 | -1.150 | 4 | 55.603 | 5.738 | 1 | 65.531 | 22.963 | 10 | 68.407 | 25.494 |
| 0.5 | 10 | 79.240 | 49.565 | 10 | 85.427 | 17.590 | 10 | 84.435 | 16.764 | 4 | 63.600 | 6.350 | 10 | 60.735 | -3.232 | 4 | 58.242 | 1.308 | 1 | 71.716 | 20.168 | 10 | 73.232 | 20.669 |
| 1 | 10 | 96.341 | 41.699 | 10 | 98.573 | 22.939 | 10 | 89.427 | 11.507 | 4 | 68.104 | -4.141 | 10 | 71.956 | -4.551 | 4 | 67.398 | -2.423 | 1 | 75.050 | 5.152 | 10 | 86.453 | 17.679 |
| 2 | 10 | 95.327 | 27.275 | 10 | 98.341 | 17.259 | 10 | 97.248 | 14.256 | 4 | 81.427 | 3.535 | 10 | 86.499 | 6.667 | 4 | 74.918 | -5.456 | 1 | 77.515 | 2.026 | 10 | 93.251 | 21.896 |
| 4 | 10 | 96.684 | 21.920 | 10 | 99.032 | 12.414 | 10 | 97.748 | 10.678 | 4 | 94.204 | 10.518 | 10 | 87.687 | 1.565 | 4 | 88.680 | 6.030 | 1 | 90.829 | 11.465 | 10 | 90.348 | 9.691 |
| 0 | 20 | 21.407 | 0.000 | 20 | 36.002 | 0.000 | 20 | 73.404 | 0.000 | 8 | 95.089 | 0.000 | 20 | 81.117 | 0.000 | 8 | 42.886 | 0.000 | 2 | 56.971 | 0.000 | 20 | 53.562 | 0.000 |
| 0.25 | 20 | 62.069 | 39.505 | 20 | 69.170 | 32.419 | 20 | 87.687 | 13.971 | 8 | 96.157 | 0.627 | 20 | 80.203 | -3.828 | 8 | 46.894 | 0.260 | 2 | 67.613 | 20.838 | 20 | 65.059 | 21.793 |
| 0.5 | 20 | 77.167 | 48.745 | 20 | 75.569 | 25.544 | 20 | 90.057 | 12.074 | 8 | 97.385 | 0.817 | 20 | 78.600 | -8.563 | 8 | 52.239 | -1.558 | 2 | 70.145 | 13.887 | 20 | 76.588 | 25.946 |
| 1 | 20 | 96.467 | 42.860 | 20 | 84.078 | 19.780 | 20 | 98.019 | 13.983 | 8 | 98.230 | 0.474 | 20 | 84.848 | -6.806 | 8 | 66.774 | -0.066 | 2 | 74.280 | 1.341 | 20 | 94.787 | 29.466 |
| 2 | 20 | 96.038 | 28.837 | 20 | 95.386 | 24.600 | 20 | 98.825 | 10.481 | 8 | 98.379 | 0.146 | 20 | 89.577 | -3.365 | 8 | 76.395 | -1.594 | 2 | 85.310 | 10.284 | 20 | 92.585 | 21.378 |
| 4 | 20 | 97.717 | 23.711 | 20 | 97.539 | 18.287 | 20 | 99.039 | 7.907 | 8 | 98.581 | -0.138 | 20 | 90.286 | -4.971 | 8 | 95.708 | 16.139 | 2 | 93.176 | 12.949 | 20 | 92.778 | 13.282 |
| **(E) RCC4-VHL** | | | | | | | | | | | | | | | | | | | | | | | | |
|  | **Bis:Len** | | | **Bis:Paz** | | | **Bis:Cabo** | | | **Bis:Suni** | | | **Bis:Soraf** | | | **Bis:Axit** | | | **Bis:Tem** | | | **Bis:Ever** | | |
| Bis (µM) | Len  (µM) | RI | SS | Paz  (µM) | RI | SS | Cabo (µM) | RI | SS | Suni (µM) | RI | SS | Soraf (µM) | RI | SS | Axit (µM) | RI | SS | Tem (µM) | RI | SS | Ever (µM) | RI | SS |
| 0 | 0 | 0.000 | 0.000 | 0 | 0.000 | 0.000 | 0 | 0.000 | 0.000 | 0 | 0.000 | 0.000 | 0 | 0.000 | 0.000 | 0 | 0.000 | 0.000 | 0 | 0.000 | 0.000 | 0 | 0.000 | 0.000 |
| 0.25 | 0 | 9.885 | 0.000 | 0 | 16.796 | 0.000 | 0 | 12.119 | 0.000 | 0 | 6.253 | 0.000 | 0 | 8.149 | 0.000 | 0 | 15.811 | 0.000 | 0 | 5.642 | 0.000 | 0 | 6.240 | 0.000 |
| 0.5 | 0 | 22.029 | 0.000 | 0 | 29.163 | 0.000 | 0 | 19.598 | 0.000 | 0 | 19.088 | 0.000 | 0 | 20.318 | 0.000 | 0 | 23.424 | 0.000 | 0 | 13.170 | 0.000 | 0 | 18.475 | 0.000 |
| 1 | 0 | 40.106 | 0.000 | 0 | 48.619 | 0.000 | 0 | 42.191 | 0.000 | 0 | 36.065 | 0.000 | 0 | 38.617 | 0.000 | 0 | 42.161 | 0.000 | 0 | 35.659 | 0.000 | 0 | 38.961 | 0.000 |
| 2 | 0 | 62.355 | 0.000 | 0 | 66.293 | 0.000 | 0 | 61.035 | 0.000 | 0 | 57.539 | 0.000 | 0 | 57.992 | 0.000 | 0 | 64.222 | 0.000 | 0 | 55.790 | 0.000 | 0 | 58.019 | 0.000 |
| 4 | 0 | 70.876 | 0.000 | 0 | 75.350 | 0.000 | 0 | 73.474 | 0.000 | 0 | 66.783 | 0.000 | 0 | 68.955 | 0.000 | 0 | 73.756 | 0.000 | 0 | 66.915 | 0.000 | 0 | 67.841 | 0.000 |
| 0 | 1.25 | 9.902 | 0.000 | 1.25 | 15.744 | 0.000 | 1.25 | 16.572 | 0.000 | 0.5 | 14.957 | 0.000 | 1.25 | 10.522 | 0.000 | 0.5 | 14.806 | 0.000 | 0.125 | 50.203 | 0.000 | 1.25 | 52.343 | 0.000 |
| 0.25 | 1.25 | 31.834 | 23.523 | 1.25 | 36.959 | 21.503 | 1.25 | 34.416 | 21.042 | 0.5 | 23.827 | 13.657 | 1.25 | 19.840 | 11.708 | 0.5 | 31.095 | 16.105 | 0.125 | 55.248 | 18.946 | 1.25 | 63.243 | 27.180 |
| 0.5 | 1.25 | 51.529 | 31.938 | 1.25 | 50.639 | 23.321 | 1.25 | 42.500 | 22.115 | 0.5 | 37.345 | 15.216 | 1.25 | 31.060 | 10.819 | 0.5 | 32.566 | 9.152 | 0.125 | 59.055 | 17.734 | 1.25 | 66.786 | 22.011 |
| 1 | 1.25 | 76.773 | 39.974 | 1.25 | 66.552 | 19.656 | 1.25 | 63.521 | 21.470 | 0.5 | 50.733 | 12.292 | 1.25 | 47.408 | 8.904 | 0.5 | 47.394 | 4.980 | 0.125 | 62.272 | 3.759 | 1.25 | 70.985 | 11.118 |
| 2 | 1.25 | 88.489 | 28.529 | 1.25 | 75.706 | 10.159 | 1.25 | 72.628 | 11.284 | 0.5 | 62.963 | 3.371 | 1.25 | 64.758 | 6.922 | 0.5 | 62.869 | -2.221 | 0.125 | 68.257 | -4.808 | 1.25 | 71.890 | -2.897 |
| 4 | 1.25 | 88.457 | 19.164 | 1.25 | 82.156 | 7.339 | 1.25 | 79.666 | 5.734 | 0.5 | 66.154 | -2.769 | 1.25 | 69.366 | 0.048 | 0.5 | 72.909 | -1.461 | 0.125 | 71.245 | -9.947 | 1.25 | 80.479 | 0.383 |
| 0 | 2.5 | 15.758 | 0.000 | 2.5 | 29.523 | 0.000 | 2.5 | 26.926 | 0.000 | 1 | 27.898 | 0.000 | 2.5 | 16.147 | 0.000 | 1 | 26.260 | 0.000 | 0.25 | 52.642 | 0.000 | 2.5 | 54.879 | 0.000 |
| 0.25 | 2.5 | 40.481 | 26.643 | 2.5 | 54.064 | 25.890 | 2.5 | 44.306 | 20.435 | 1 | 39.976 | 16.638 | 2.5 | 26.702 | 12.986 | 1 | 37.178 | 10.267 | 0.25 | 57.750 | 18.411 | 2.5 | 66.437 | 27.334 |
| 0.5 | 2.5 | 62.822 | 38.839 | 2.5 | 67.993 | 30.301 | 2.5 | 57.351 | 28.206 | 1 | 50.783 | 17.233 | 2.5 | 36.678 | 11.558 | 1 | 42.650 | 9.097 | 0.25 | 62.055 | 18.128 | 2.5 | 70.377 | 23.197 |
| 1 | 2.5 | 84.196 | 43.895 | 2.5 | 75.699 | 20.717 | 2.5 | 70.716 | 21.862 | 1 | 59.603 | 11.945 | 2.5 | 51.965 | 9.727 | 1 | 53.353 | 3.026 | 0.25 | 66.521 | 6.624 | 2.5 | 72.087 | 9.955 |
| 2 | 2.5 | 87.917 | 25.220 | 2.5 | 81.003 | 10.037 | 2.5 | 75.798 | 9.623 | 1 | 65.692 | -0.368 | 2.5 | 67.104 | 6.631 | 1 | 65.576 | -4.562 | 0.25 | 68.179 | -6.646 | 2.5 | 75.969 | 0.597 |
| 4 | 2.5 | 89.869 | 18.648 | 2.5 | 90.572 | 12.532 | 2.5 | 86.994 | 10.525 | 1 | 68.731 | -5.204 | 2.5 | 72.283 | 1.139 | 1 | 71.734 | -6.829 | 0.25 | 71.373 | -11.085 | 2.5 | 85.562 | 5.591 |
| 0 | 5 | 21.752 | 0.000 | 5 | 48.841 | 0.000 | 5 | 38.052 | 0.000 | 2 | 47.334 | 0.000 | 5 | 22.884 | 0.000 | 2 | 38.307 | 0.000 | 0.5 | 51.863 | 0.000 | 5 | 53.771 | 0.000 |
| 0.25 | 5 | 46.654 | 26.883 | 5 | 73.036 | 26.215 | 5 | 64.956 | 31.260 | 2 | 55.057 | 10.932 | 5 | 34.242 | 13.765 | 2 | 45.789 | 6.680 | 0.5 | 59.459 | 21.767 | 5 | 69.749 | 33.352 |
| 0.5 | 5 | 68.191 | 39.078 | 5 | 79.018 | 24.615 | 5 | 72.087 | 33.349 | 2 | 63.425 | 11.893 | 5 | 46.370 | 15.714 | 2 | 49.132 | 4.276 | 0.5 | 63.280 | 20.775 | 5 | 73.416 | 28.637 |
| 1 | 5 | 88.499 | 44.275 | 5 | 81.797 | 14.383 | 5 | 77.090 | 20.725 | 2 | 68.868 | 6.923 | 5 | 60.457 | 14.061 | 2 | 57.493 | -1.490 | 0.5 | 67.560 | 8.752 | 5 | 75.000 | 14.852 |
| 2 | 5 | 87.772 | 22.317 | 5 | 92.581 | 14.696 | 5 | 84.599 | 14.017 | 2 | 69.673 | -6.114 | 5 | 70.153 | 6.547 | 2 | 66.750 | -8.957 | 0.5 | 69.617 | -4.264 | 5 | 82.232 | 9.473 |
| 4 | 5 | 90.573 | 17.302 | 5 | 92.547 | 8.389 | 5 | 91.924 | 12.300 | 2 | 76.322 | -4.731 | 5 | 79.045 | 6.056 | 2 | 73.863 | -8.588 | 0.5 | 72.688 | -9.000 | 5 | 86.680 | 7.632 |
| 0 | 10 | 29.300 | 0.000 | 10 | 67.894 | 0.000 | 10 | 62.470 | 0.000 | 4 | 58.307 | 0.000 | 10 | 60.904 | 0.000 | 4 | 51.057 | 0.000 | 1 | 51.498 | 0.000 | 10 | 54.368 | 0.000 |
| 0.25 | 10 | 55.147 | 27.980 | 10 | 82.430 | 15.698 | 10 | 78.783 | 18.954 | 4 | 63.536 | 7.672 | 10 | 62.025 | 1.857 | 4 | 56.203 | 4.387 | 1 | 58.244 | 20.779 | 10 | 70.328 | 33.166 |
| 0.5 | 10 | 71.474 | 35.546 | 10 | 85.028 | 13.349 | 10 | 79.294 | 15.858 | 4 | 68.970 | 7.112 | 10 | 66.857 | 1.391 | 4 | 57.955 | 1.440 | 1 | 64.288 | 22.566 | 10 | 71.439 | 25.268 |
| 1 | 10 | 90.799 | 41.317 | 10 | 93.193 | 14.396 | 10 | 85.210 | 11.514 | 4 | 72.586 | 2.407 | 10 | 70.160 | -3.686 | 4 | 62.041 | -6.075 | 1 | 66.266 | 7.488 | 10 | 79.503 | 20.065 |
| 2 | 10 | 87.241 | 18.279 | 10 | 95.346 | 9.225 | 10 | 92.604 | 10.707 | 4 | 75.267 | -5.645 | 10 | 74.818 | -7.780 | 4 | 70.058 | -11.222 | 1 | 69.225 | -4.502 | 10 | 87.550 | 15.930 |
| 4 | 10 | 91.115 | 15.229 | 10 | 94.928 | 4.806 | 10 | 90.639 | 2.335 | 4 | 84.966 | 0.372 | 10 | 83.144 | -3.812 | 4 | 77.910 | -8.378 | 1 | 77.891 | -2.198 | 10 | 85.415 | 5.677 |
| 0 | 20 | 27.935 | 0.000 | 20 | 36.033 | 0.000 | 20 | 80.555 | 0.000 | 8 | 83.280 | 0.000 | 20 | 80.914 | 0.000 | 8 | 51.221 | 0.000 | 2 | 48.773 | 0.000 | 20 | 52.693 | 0.000 |
| 0.25 | 20 | 48.769 | 22.444 | 20 | 67.660 | 34.376 | 20 | 84.725 | 4.916 | 8 | 85.130 | 2.803 | 20 | 79.494 | -1.266 | 8 | 56.133 | 4.123 | 2 | 60.180 | 27.384 | 20 | 66.669 | 31.060 |
| 0.5 | 20 | 69.353 | 34.501 | 20 | 73.277 | 30.209 | 20 | 87.130 | 5.764 | 8 | 87.156 | 2.408 | 20 | 79.663 | -3.911 | 8 | 58.156 | 1.506 | 2 | 64.983 | 27.264 | 20 | 71.226 | 27.274 |
| 1 | 20 | 91.286 | 42.847 | 20 | 77.625 | 18.433 | 20 | 94.912 | 8.914 | 8 | 90.942 | 3.114 | 20 | 80.944 | -6.756 | 8 | 64.469 | -3.397 | 2 | 66.348 | 10.419 | 20 | 88.975 | 34.021 |
| 2 | 20 | 87.661 | 19.367 | 20 | 90.381 | 17.976 | 20 | 93.295 | 2.257 | 8 | 95.702 | 3.974 | 20 | 85.090 | -6.687 | 8 | 71.408 | -9.741 | 2 | 74.419 | 4.037 | 20 | 85.816 | 14.863 |
| 4 | 20 | 91.544 | 16.185 | 20 | 93.020 | 13.209 | 20 | 94.256 | 0.186 | 8 | 94.700 | 0.959 | 20 | 88.441 | -5.539 | 8 | 86.263 | 1.213 | 2 | 86.466 | 10.146 | 20 | 87.365 | 9.099 |

Bis: Bisantrene; Len: Lenvatinib; Paz: Pazopanib; Cabo: Cabozantinib; Suni: Sunitinib; Soraf: Sorafenib; Axit: Axitinib; Tem: Temsirolimus; Ever: Everolimus; RI: Relative inhibition; SS: Synergy Score.

Supplementary Figure 1: Cytotoxicity of single agent ccRCC drugs in 786-O, Caki-1, Caki-2, RCC4-EV and RCC4-VHL ccRCC cells treated for 72 h with the indicated concentrations of: (A) lenvatinib; (B) pazopanib; (C) cabozantinib; (D) sunitinib; (E) sorafenib; (F) axitinib; (G) temsirolimus; and (H) everolimus. Cell viability was determined using a resazurin metabolic assay and is expressed as a percentage of untreated control cells. Mean ± SEM, n=3.

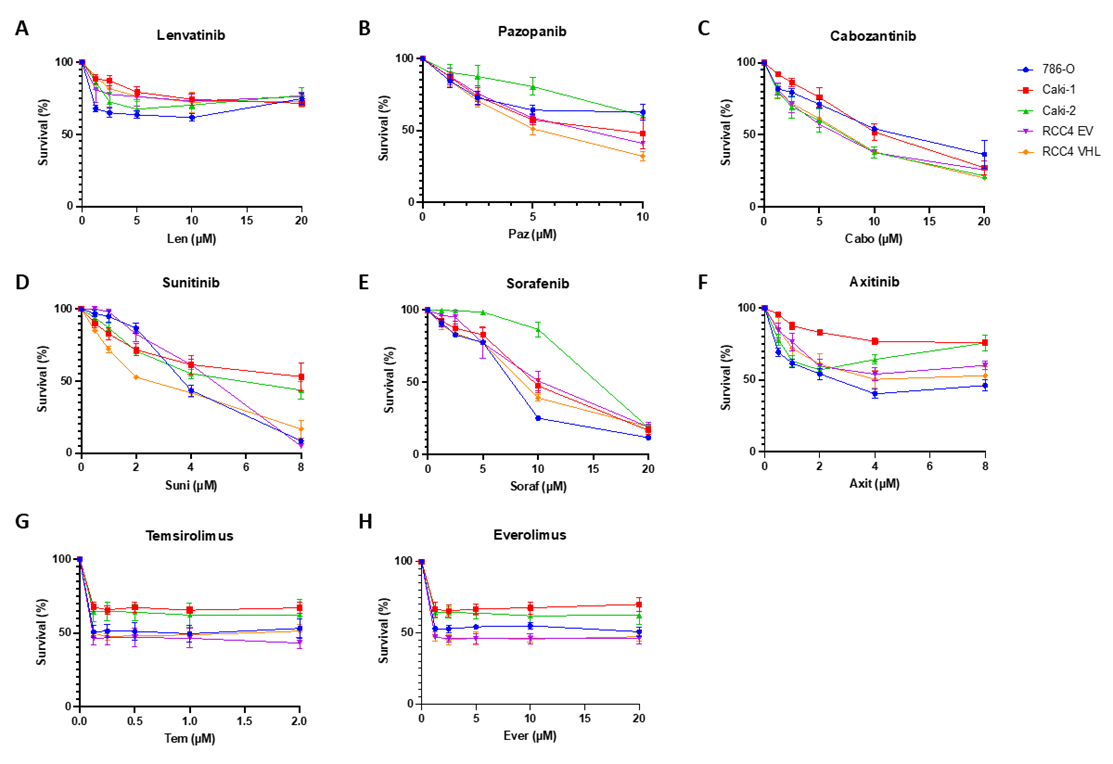

**Supplementary Figure 2: Cell viability in response to combined treatment with bisantrene and ccRCC drugs in ccRCC cell lines.** ccRCC cell lines 786-O, Caki-1, Caki-2, RCC-EV and RCC-VHL were treated with indicated concentrations of bisantrene and ccRCC drugs for 72 h and cell viability determined using a resazurin assay. ccRCC drugs used: **(A)** sunitinib**; (B)** sorafenib; **(C)** axitinib; **(D)** temsirolimus; and **(E)** everolimus. Average percent viability relative to untreated cells is presented as a heat map, where blue indicates high viability and red indicates low viability. + indicates synergy, as assessed by the fractional product method of Webb (Webb JL. Enzyme and Metabolic Inhibitors: Academic Press1963).

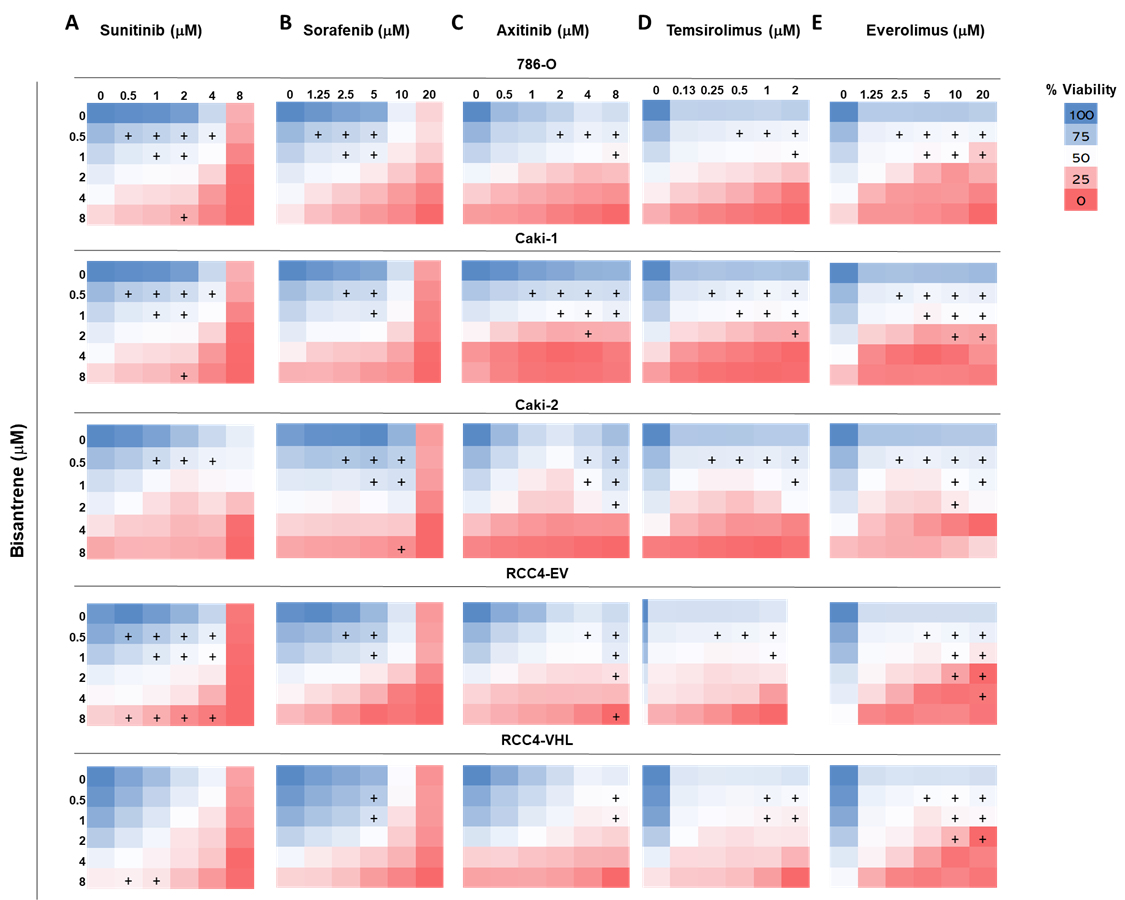

**Supplementary Figure 3:** 2D and 3D visualization of predicted Bliss scores in: **(A)** 786-O, **(B)** Caki-1, **(C)** Caki-2, **(D)** RCC4-EV, and **(E)** RCC$-VHL cells at each concentration point, with red to green scale indicating areas of synergy to antagonism, and the average synergy score. The most synergistic 2x2 area is indicated with a white box. **(i)** lenvatinib; **(ii)** pazopanib; **(iii)** cabozantinib; **(iv)** sunitinib**; (v)** sorafenib; **(vi)** axitinib; **(vii)** temsirolimus; and **(viii)** everolimus.

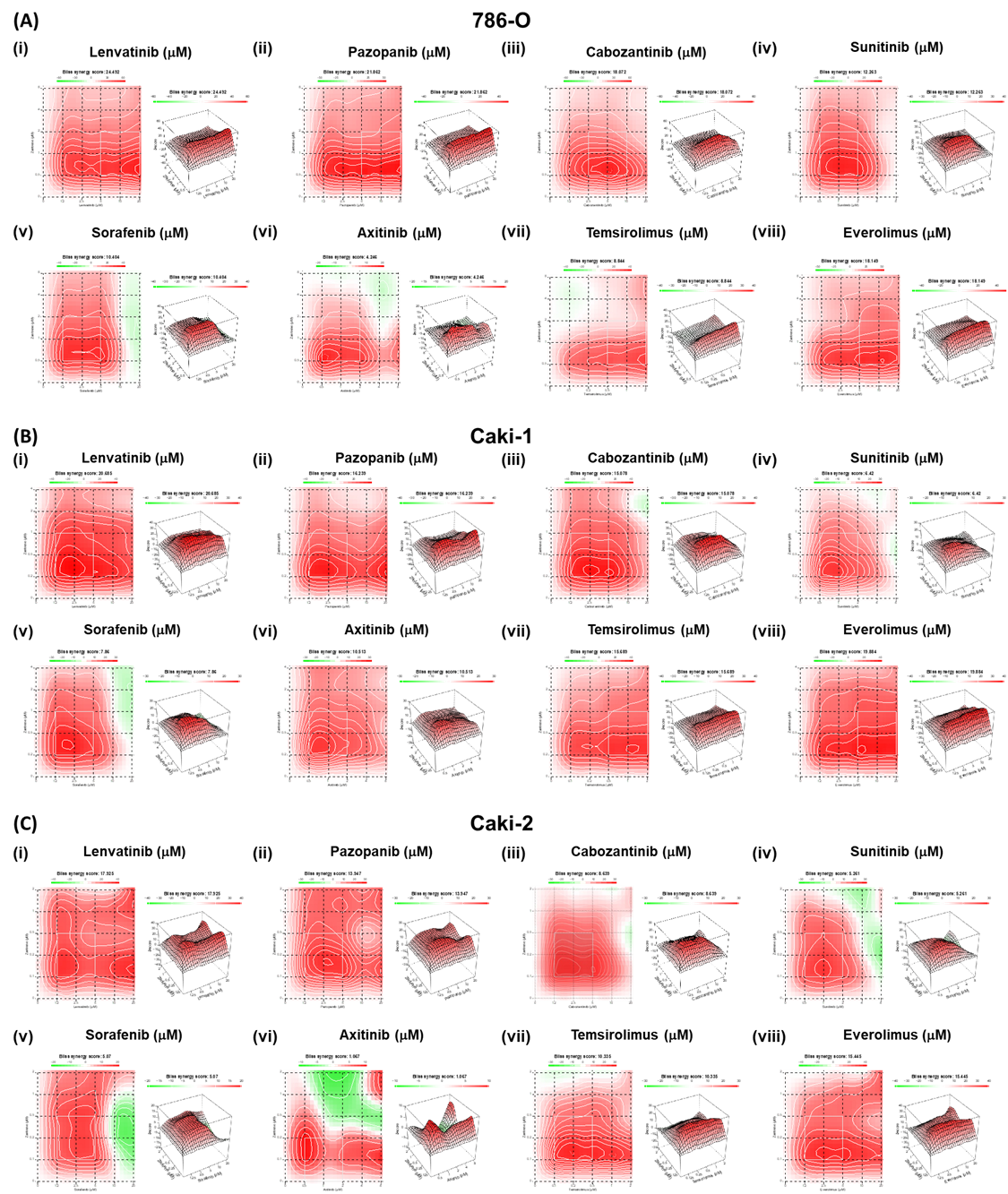

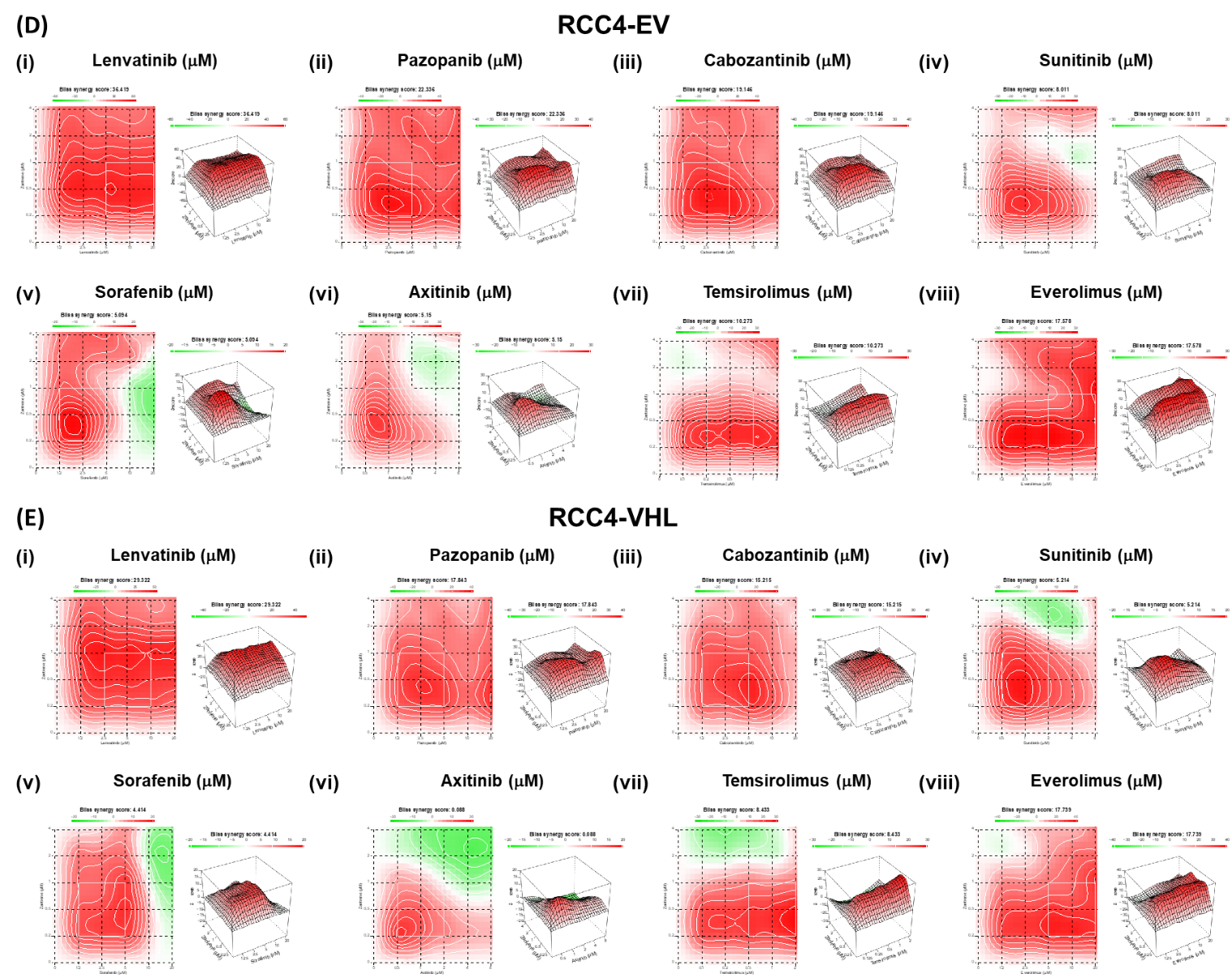

**Supplementary Figure 4: (A)** Cytotoxicity of single agent tivozanib in 786-O cells. Cells were treated for 72 h with the indicated drug concentrations and cell viability was determined using a resazurin metabolic assay. Cell viability is expressed as a percentage of untreated control cells. **(B)** Cell viability in response to combined drug treatment with bisantrene and tivozanib in 786-O cells. 786-O cells were treated with the indicated concentrations of bisantrene and tivozanib for 72 h and cell viability determined using a resazurin assay. Average percent viability relative to untreated cells is presented as a heat map, where blue indicates high viability and red indicates low viability. + signs indicate synergy, as assessed by the fractional product method of Webb. **(C)** Bliss synergy analysis of bisantrene and tivozanib in 786-O cells. 2D and 3D visualization of predicted Bliss scores in 786-O cells at each concentration point, with red to green scale indicating areas of synergy to antagonism, and the average synergy score. The most synergistic 2x2 area is indicated with a white box. OSS = Overall synergy score; MSAS = Most synergistic area score.

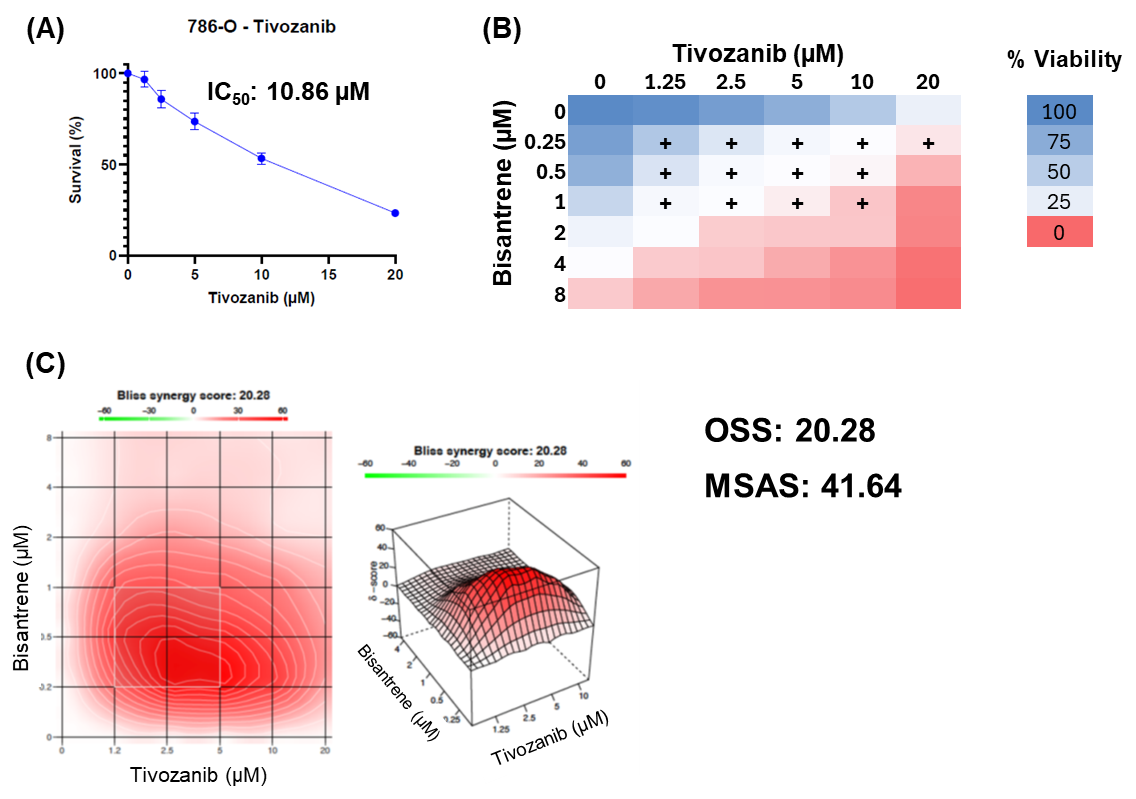
